# RNA functional control by hydrolysis reversible acylation

**DOI:** 10.1101/2024.08.29.610419

**Authors:** Kaizheng Liu, Anna M Kietrys

## Abstract

Reversible 2′-OH acylation is a powerful strategy for switching RNA function, but existing systems often rely on nonphysiological or cytotoxic triggers for deacylation. Here we present **EST1A**, a hydrolysis-responsive 2′-OH acylating reagent whose RNA adducts are efficiently removed by endogenous esterases *in vitro* and *in cellulo*. **EST1A** acylates model oligonucleotides, an EGFP-targeting antisense strand, and reporter mRNAs, thereby modulating their activity; notably, the acylated antisense strand shows enhanced EGFP knockdown in HepG2 cells. By tuning carboxylesterase and cholinesterase activity and comparing **EST1A-**acylated mCherry mRNA across noncancerous and cancer-derived cell lines, we reveal a positive correlation between intracellular esterase activity and functional recovery of acylated RNA. These results establish **EST1A**-mediated, hydrolysis-responsive 2′-OH acylation as a simple platform for enzyme-guided, cell-selective activation of RNA function and point toward esterase-activated RNA therapeutics.

**Entry for the Table of Contents:** Liu et al. introduce **EST1A**, a hydrolysis-responsive 2′-OH acylating reagent whose RNA adducts are removed by endogenous esterases or histidine, enabling reversible control of RNA function. By exploiting differences in esterase activity between noncancerous and cancer-derived cell lines, **EST1A**-treated mRNA exhibits enzyme-guided and cell-selective translational reactivation.

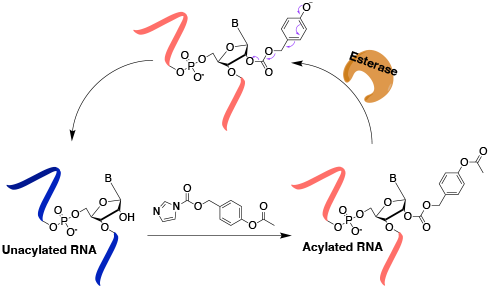

---

RNA is a multifunctional biomacromolecule that participates in diverse biochemical processes, including protein synthesis, gene expression regulation, and RNA modification.^[1]^ Owing to this versatility, RNA has become indispensable in both basic research and therapeutic applications.^[2]^ The great potential of RNA in these contexts has driven the development of methods for activity control, interrogation of biological roles, and labeling.^[3]^ Previous studies have demonstrated that reversible 2′-OH acylation is an effective strategy for modulating RNA function.^[4]^ Acylation transiently deactivates RNAs (for example, preventing mRNA translation), whereas subsequent deacylation restores their activity.^[4c]^ Moreover, as a post-synthetic modification, 2′-OH acylation can be applied not only to short oligonucleotides but also to long and structured RNAs.^[5]^ However, currently available deacylation strategies often rely on toxic small molecules or UV irradiation, substantially limiting their therapeutic potential.^[4c, 4d, 6]^

To eliminate cytotoxicity at the deacylation step and better explore the therapeutic potential of acylation, we envisioned that endogenous enzymes could serve as biocompatible deacylating reagents, as they naturally exist in cells. Moreover, Distinct cell lines display characteristic enzyme profiles, suggesting a route to cell-selective prodrug activation. In this work, we focused on esterases, which cleave ester bonds and are closely linked to cellular energy metabolism.^[7]^ Carboxylesterases (CESs) are also frequently overexpressed in cancer and have been implicated in tumor migration.^[8]^ Beyond CESs, we also considered other esterase families that might contribute to deacylation. Cholinesterases (ChEs) are best known as membrane-associated enzymes that terminate neurotransmission, but their presence in the cytoplasm of neuronal cells, especially under stress, has also been reported. ^[9]^ In addition to these enzyme-mediated pathways, we found that simple nucleophiles such as imidazole and histidine can promote ester hydrolysis. Although the intracellular concentration of free histidine is relatively low, this nonenzymatic pathway is still useful for *in vitro* applications. To enable hydrolysis-mediated deacylation, we incorporated a hydrolysis-responsive ester linkage into the acylating reagent. As this linkage is readily cleaved in the intracellular environment, we anticipated gradual restoration of RNA activity via deacylation *in cellulo*.^[10]^ Since the deacylation (restoration) rate depends on CES expression levels, this strategy inherently enables cell-specific activation of RNA-based drugs. Accordingly, we prepared two acylating reagents, **EST1A** and **EST1B**, to implement this reversible strategy (Figure 1A). Guided by prior 2′-OH acylation studies, imidazole and 2-chloroimidazole were chosen as leaving groups.^[4b, 4d]^ A benzyl acetate unit was incorporated as the hydrolysis-responsive trigger. Upon ester cleavage, the resulting benzyl phenol intermediate undergoes a 1,6-elimination; subsequent release of a quinone methide and carbon dioxide (Figure 1B) regenerates the free 2′-OH.

**Figure 1.**
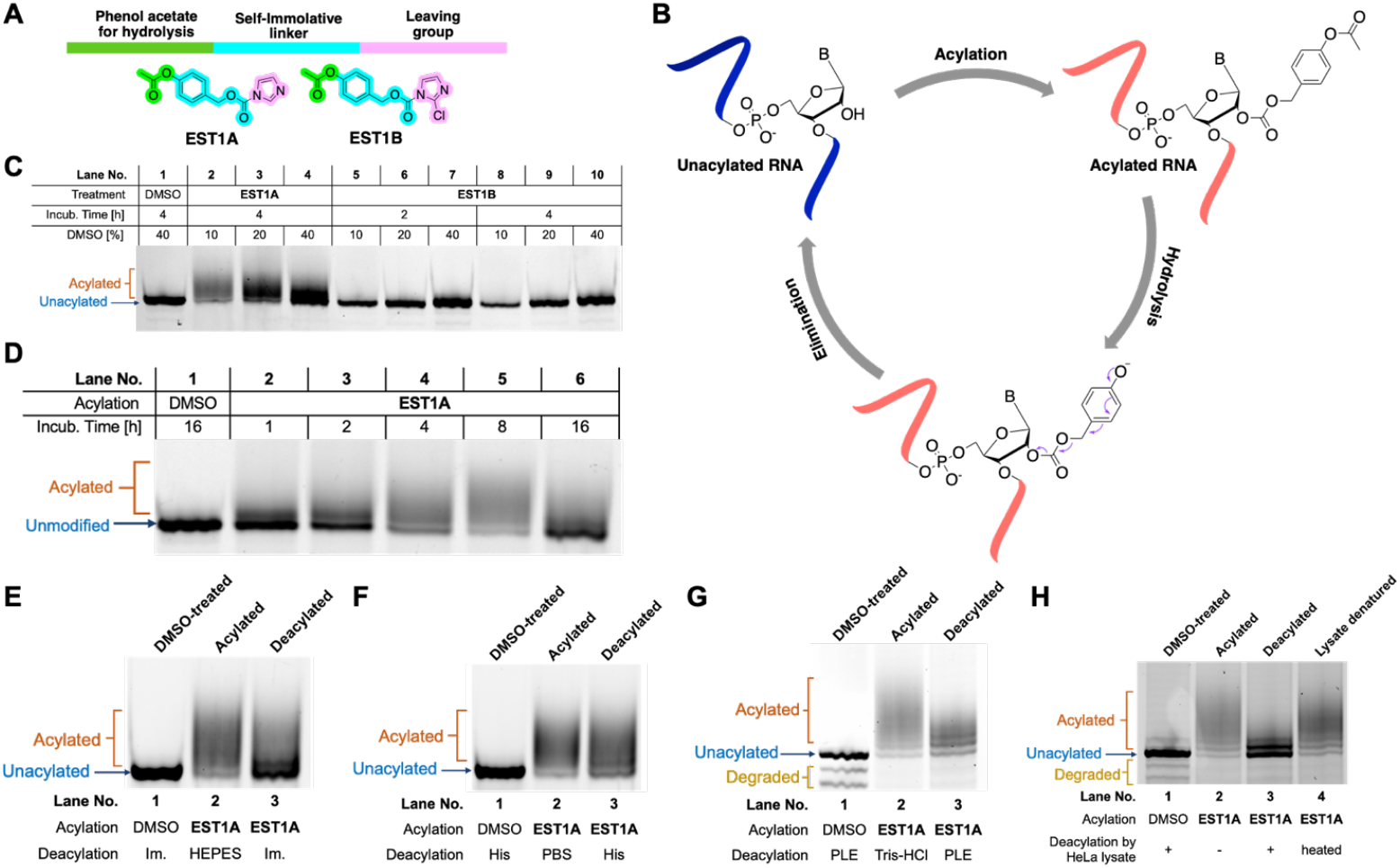
The structure, mechanism, and reactivity screening of acylation reagents. (A) Design principle of EST1A and EST1B. (B) The mechanism of acylation removal by esterase. (C) Acylation activity of **EST1A** and **EST1B**. 120 nM Cy3-U_16_ acylated by 100 mM **EST1A** or **EST1B** at 37 °C for 4 hours. The gel was scanned in the Cy3 channel. (D) Acylation of Cy3-U_16_ RNA by 100 mM **EST1A** in 10% DMSO at 37 °C for 2-16 hours. (E and F) PAGE analysis: deacylation by imidazole (D) and histidine (E). Acylation: 120 nM Cy3-U_16_ acylated by 100 mM **EST1A** at 37 °C for 6 hours. Deacylation: 120 nM acylated Cy3-U_16_ RNA incubated with 50 mM imidazole in 50 mM HEPES or 50 mM histidine in PBS at 37 °C for 4 hours. The gel was scanned in the Cy3 channel. (G and H) PAGE analysis: deacylation by porcine liver esterase (D) and HeLa cell lysate. Acylation: 5 µM antisense strand of EGFP siRNA incubated with 50 mM **EST1A** in 10% DMSO at 37 °C for 4 hours. Deacylation: 5 µM acylated antisense strand incubated with 40 U/mL PLE or HeLa cell lysate (total protein: 0.5 µg/µL) in 50 mM Tris-HCl (pH 8.0) at 37 °C for 4 hours. The gel was scanned in the SYBR-gold channel. Im. – imidazole; His – histidine.

We first assessed the reactivity of **EST1A** and **EST1B** using a 5′-Cy3-labeled 16-mer polyuridine (Cy3-U_16_). 120 nM Cy3-U16 RNA was treated with 100 mM **EST1A** in different percentage of DMSO at 37 °C for 4 hours, followed by analysis by 30% denaturing polyacrylamide gel electrophoresis (30% dPAGE, also see Figure S1-2). In Figure 1C, lanes 2–4 showed slower-migrating bands relative to the unmodified control, consistent with formation of acylation adducts by **EST1A**. In contrast, **EST1B**, although bearing a more activated leaving group, did not generate detectable products under any of the solvent conditions tested (lanes 5–10 in Figure 1C, and Figure S1), likely due to limited solubility or rapid hydrolysis.^[4b]^ We next examined esterase-triggered removal of acylation. During the optimization of acylating conditions of **EST1A**, prolonging the incubation time led to lower levels of acylation (Figure 1D and S3). This unexpected observation can be attributed to the catalytical activity of imidazole (generated during acylation) in the ester hydrolysis, making it a potential deacylating reagent.^[11]^ To test this, we prepared acylated Cy3-U16 and treated it with imidazole. PAGE analysis showed the disappearance of the slower-migrating, up-shifted band, indicating efficient deacylation by imidazole (lane 3 in Figure 1E). We next examined histidine, an amino acid that bears an imidazole side chain and serves as the catalytic base in esterase active sites.^[12]^ Consistent with our hypothesis, histidine also promoted reversal of acylation (Figure 1F and S4). To minimize degradation by RNases, we used antisense strand of EGFP siRNA (siEGFP-AS) bearing a dTdT overhang as the substrate. This strand was treated with 50 mM **EST1A** in 10% DMSO at 37 °C for 4 h, purified by ethanol precipitation, and then incubated with either porcine liver esterase (PLE) or Tris–HCl buffer. In 30% dPAGE image (Figure 1G and S5), the band in lane 3 (deacylated sample) shifted toward the position of unmodified RNA relative to lane 2 (acylated sample), indicating substantial removal of acylation adducts (see Figure S7 for MALDI-TOF). Deacylation was next evaluated under more physiologically relevant conditions using HeLa cell lysate. Incubation with native lysate at 37 °C for 4 h led to fading of the upshifted bands corresponding to acylated RNA strands (lane 3, Figure 1H), whereas denatured lysate (heated at 95 °C for 5 min) failed to restore the species with low mobility, indicating that enzymatic activity was required for acylation reversal.

After validating the reversibility of **EST1A** using short oligonucleotides, we next examined whether it could control the folding of longer RNAs. As a model, we selected the F-30 broccoli aptamer. Properly folded F-30 broccoli binds DFHBI, and the resulting aptamer–DFHBI complex emits fluorescence by restricting intramolecular motion of the dye. Acylation of the aptamer was expected to interfere with its folding and thereby disrupt this interaction (Figure 2A). We incubated F-30 broccoli aptamer with DMSO or increasing concentrations of **EST1A** and folded it in HEPES folding buffer and then mixed with DFHBI (20 equiv.). Aptamers treated with higher concentrations of **EST1A** gave weaker fluorescence signals (Figure 2B), confirming that **EST1A** disrupts the aptamer–DFHBI interaction. We then evaluated deacylation by imidazole and histidine. Imidazole was more effective: treatment of acylated aptamer with 50 mM imidazole in HEPES buffer (pH 7.5) restored 61% relative fluorescence, compared with 32% for the corresponding acylated control (Figure 2C). Acylated aptamer incubated with 50 mM histidine in PBS (pH 7.4) recovered 37% relative fluorescence, nearly twice that of the sample incubated without histidine (19%) (Figure 2D). The higher apparent deacylation efficiency observed with imidazole may reflect buffer-dependent differences in RNA folding: in HEPES, the acylated aptamer may adopt conformations that expose acyl groups at positions less critical for folding, allowing those modifications to be preferentially removed and thereby favoring recovery of the functional structure.

**Figure 2.**
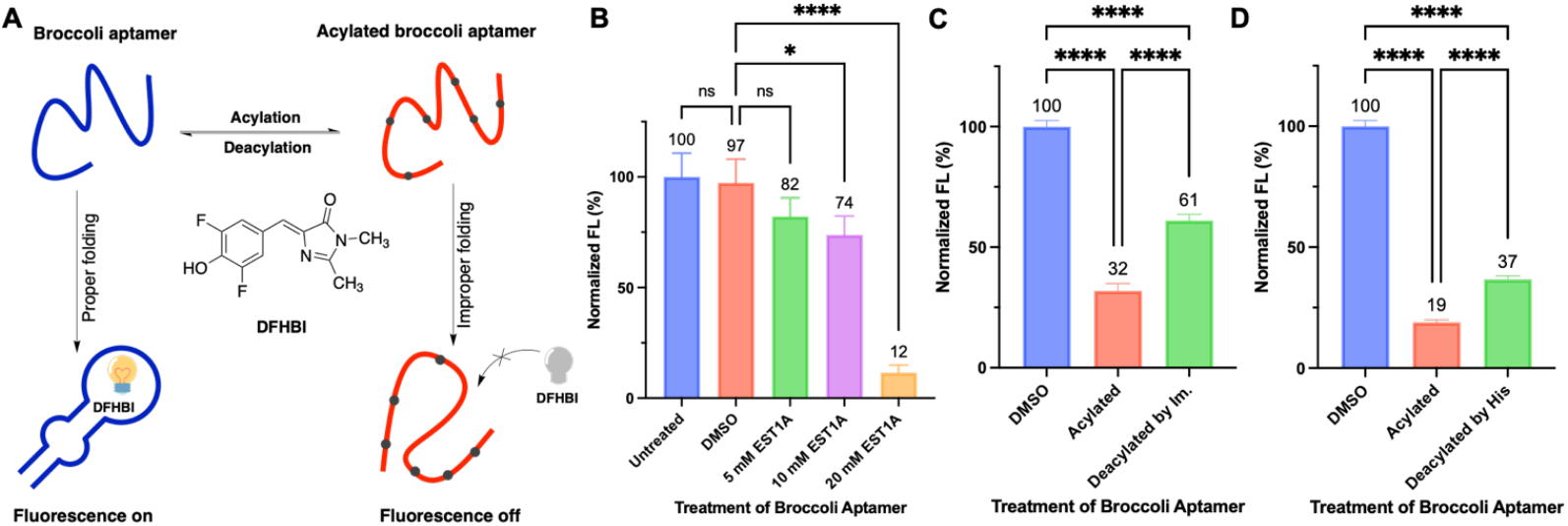
Reversible acylation of the Broccoli aptamer. (A) Schematic of **EST1A**-mediated acylation controlling the Broccoli aptamer–DFHBI interaction. (B) DFHBI fluorescence of aptamers acylated with increasing concentrations of EST1A. (C) Quantification of aptamer functional restoration by imidazole. (D) Quantification of aptamer functional restoration by histidine. Data are represented as mean ± SD (n = 3). t-test: ns, p > 0.05; *p ≤ 0.05; **p ≤ 0.01; ***p ≤ 0.001; ****p ≤ 0.0001. Acylation: 200 ng/µL aptamer incubated with 20 mM **EST1A** at 37 °C for 4 hours. Deacylation: 100 ng/µL aptamer incubated with 50 mM imidazole in 50 mM HEPES buffer or 50 mM histidine in PBS at 37 °C for 4 hours.

Next, we applied our acylating reagent to control translation of an mRNA encoding enhanced green fluorescent protein (EGFP). As previously reported, 2′-OH acylation can interfere with ribosomal reading of mRNA and thereby inhibit protein synthesis (Figure 3A). To test whether **EST1A** can modulate gene expression, we acylated 40 ng/µL EGFP mRNA with 5 mM **EST1A** at 37 °C for 4 h. After ethanol precipitation, the acylated mRNA was incubated with 50 mM histidine in PBS or 50 mM imidazole in HEPES at 37 °C for 4 h to allow deacylation. The resulting mRNAs were then transfected into HepG2 cells using Lipofectamine MessengerMax, and green fluorescence was measured after overnight incubation. Acylation strongly suppressed translation: HepG2 cells transfected with acylated EGFP mRNA showed only ~20% relative fluorescence compared with the DMSO-treated control (Figure 3B). Deacylation by histidine largely restored expression, yielding ~71% relative fluorescence (Figure 3B-C). These results show that **EST1A**-mediated reversible 2′-OH acylation can be used to regulate the translation of a long, structured mRNA in cells.

**Figure 3.**
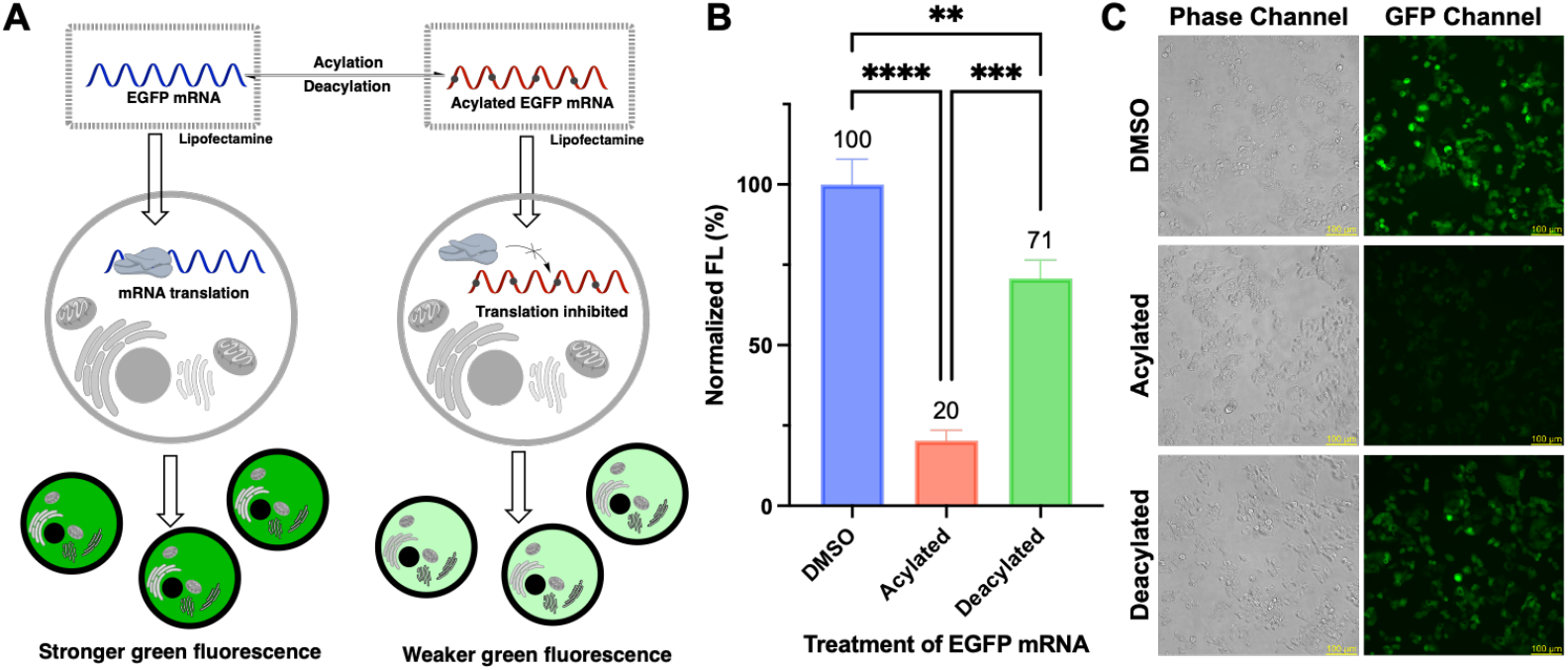
Reversible acylation of the EGFP mRNA. (A) Schematic of **EST1A**-mediated control of EGFP mRNA translation. (B) Quantification of EGFP expression following **EST1A**-mediated acylation and histidine-triggered deacylation. (C) Representative fluorescence microscopy images of HepG2 cells transfected with DMSO-treated, acylated, or deacylated EGFP mRNA. Scale bar: 100 µm. Data are represented as mean ± SD (n = 3). t-test: ns, p > 0.05; *p ≤ 0.05; **p ≤ 0.01; ***p ≤ 0.001; ****p ≤ 0.0001. Acylation: 40 ng/µL EGFP mRNA incubated with 5 mM **EST1A** at 37 °C for 4 h. Deacylation: 20 ng/µL EGFP mRNA incubated with 50 mM histidine in PBS at 37 °C for 4 h.

We then evaluated whether **EST1A**-mediated acylation modulates the activity of EGFP-targeting antisense strand in cells.^[13]^ DMSO-treated and **EST1A**-acylated siEGFP-AS were prepared and the modification was confirmed by 30% dPAGE (Figure 4A). The resulting siEGFP-AS were co-transfected with EGFP mRNA into HepG2 cells using Lipofectamine 3000 (Figure 4A).^[14]^ In contrast to mRNA and aptamer substrates, where acylation primarily suppresses activity, reversible 2′-OH acylation of siRNAs or antisense strands can enhance gene silencing by improving RNA stability, facilitating cellular uptake, and reactivating via intracellular deacylation.^[15]^ In line with this mechanism, **EST1A**-acylated antisense strand produced stronger translational suppression than the DMSO-treated one, as reflected by a larger reduction in EGFP fluorescence: 86% (DMSO-treated) versus 64% (**EST1A**-acylated) of the scrambled control (Figure 4B and 4C). To assess target silencing at the mRNA level, total RNA was isolated, converted to cDNA by reverse transcription (RT), and analyzed by quantitative PCR (qPCR). The

**Figure 4.**
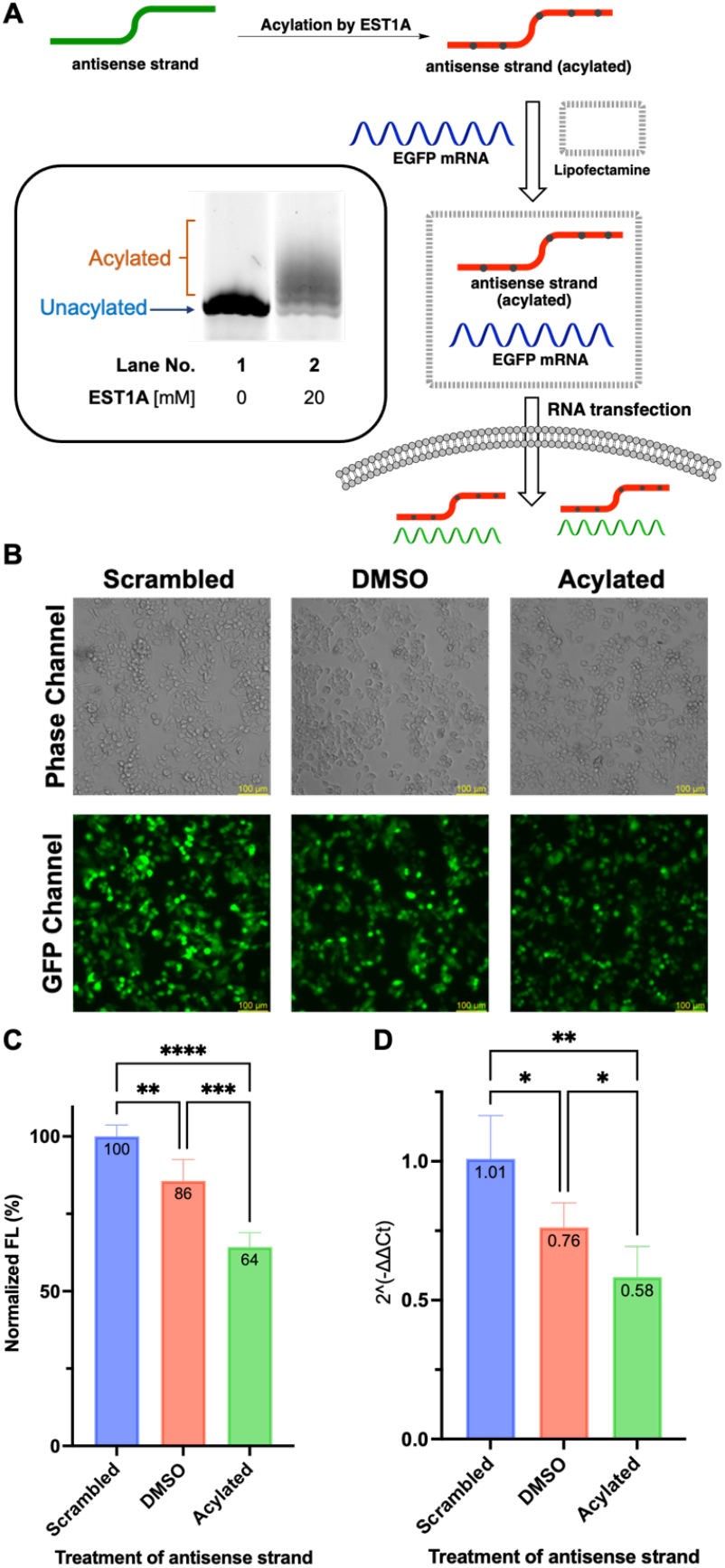
Acylation of the antisense strand of EGFP siRNA. (A) PAGE analysis of the antisense strand of EGFP siRNA (siEGFP-AS) before and after **EST1A** treatment and schematic of the co-transfection experiment. 5 µM siEGFP-AS was acylated with 20 mM **EST1A** in 10% DMSO at 37 °C for 4 h. (B) Representative fluorescence microscopy images of HepG2 cells transfected with 0.33 ng/µL EGFP mRNA and 100 nM siEGFP (scrambled control, DMSO-treated siEGFP-AS, or acylated siEGFP-AS). Scale bar: 100 µm. (C) Quantification of EGFP fluorescence intensity in HepG2 cells (mean ± SD, n = 4). (D) Relative EGFP mRNA levels quantified by RT–qPCR (mean ± SD, n = 4). t-test: ns, p > 0.05; *p ≤ 0.05; **p ≤ 0.01; ***p ≤ 0.001; ****p ≤ 0.0001.

RT-qPCR data paralleled the fluorescence readout, showing lower EGFP transcript levels in samples treated with the acylated siRNA (Figure 4D). Overall, these results indicate that **EST1A**-based reversible acylation can potentiate antisense-mediated inhibition and may be exploited to improve the performance of siRNA therapeutics.

Having established that **EST1A**-derived acylation is reversible in lysate (Figure 1H), we sought to determine whether endogenous cellular esterases could also remove the acylation groups and restore RNA function *in cellulo*. EGFP mRNA was chosen as a reporter and subjected to either DMSO-treatment (control) or **EST1A**-acylation before transfection. To modulate carboxylesterase levels, HepG2 cells were pretreated with the hepatotoxic drug acetaminophen (APAP) to reduce esterase activity or with the anticancer agent 5-fluorouracil (5-FU) to enhance it (Figure 5A).^[16]^ APAP is a widely used analgesic and antipyretic; overdose causes liver injury through oxidative and endoplasmic reticulum (ER) stress, compromising ER-resident enzymes such as carboxylesterases.^[17]^ In contrast, 5-FU induces replication stress and has been reported to upregulate CES expression in cancer cells.^[16c, 18]^ EGFP mRNAs were then delivered into HepG2 cells under these different conditions. For quantification, EGFP fluorescence from cells transfected with DMSO-treated mRNA was defined as 100%, and all other signals were expressed as a percentage of this value. As shown in Figure 5B, APAP treatment (1 mM) decreased carboxylesterase activity and attenuated *in cellulo* deacylation, reducing relative fluorescence from 34.6% to 25.8% (3 mM **EST1A**) and from 25.3% to 20.6% (5 mM **EST1A**). In the enhancement assay, HepG2 cells were pre-exposed to 80 µM 5-FU prior to mRNA delivery. As shown in Figure 5C, when EGFP mRNA was acylated with 5 mM **EST1A**, cells treated with 5-FU exhibited stronger relative fluorescence (26.2%) than untreated cells (21.9%). We next asked whether this concept extends to other esterase families and cell types. We chose U87MG cells and inhibited its cellular cholinesterase by 100 µM tacrine (Figure 5A).^[19]^ The tacrine-treated U87MG cells showed lower relative fluorescence intensity (42.2%) compared with untreated cells (65.6%) when transfected with 5 mM **EST1A**-acylated mRNA, indicating the *in cellulo* deacylation was suppressed (Figure 5D). For the enhancement assay, U87MG cells were exposed to 200 µM hydrogen peroxide for 1 h before mRNA transfection. Oxidative stress has been reported to increase cholinesterase in neuronal cells and is therefore expected to favor *in cellulo* deacylation.^[9a]^ Consistent with this expectation, cells without H_2_O_2_ treatment showed 22.8% relative fluorescence, whereas oxidative stress increased the intensity to 34.6% for mRNA acylated with 5 mM **EST1A** (Figure 5E). Together with the CES perturbation experiments in HepG2 cells, these results demonstrate that tuning intracellular esterase activity—including both carboxylesterase and cholinesterase in distinct cell types—systematically modulates the functional recovery of **EST1A**-acylated mRNA *in cellulo*.

**Figure 5.**
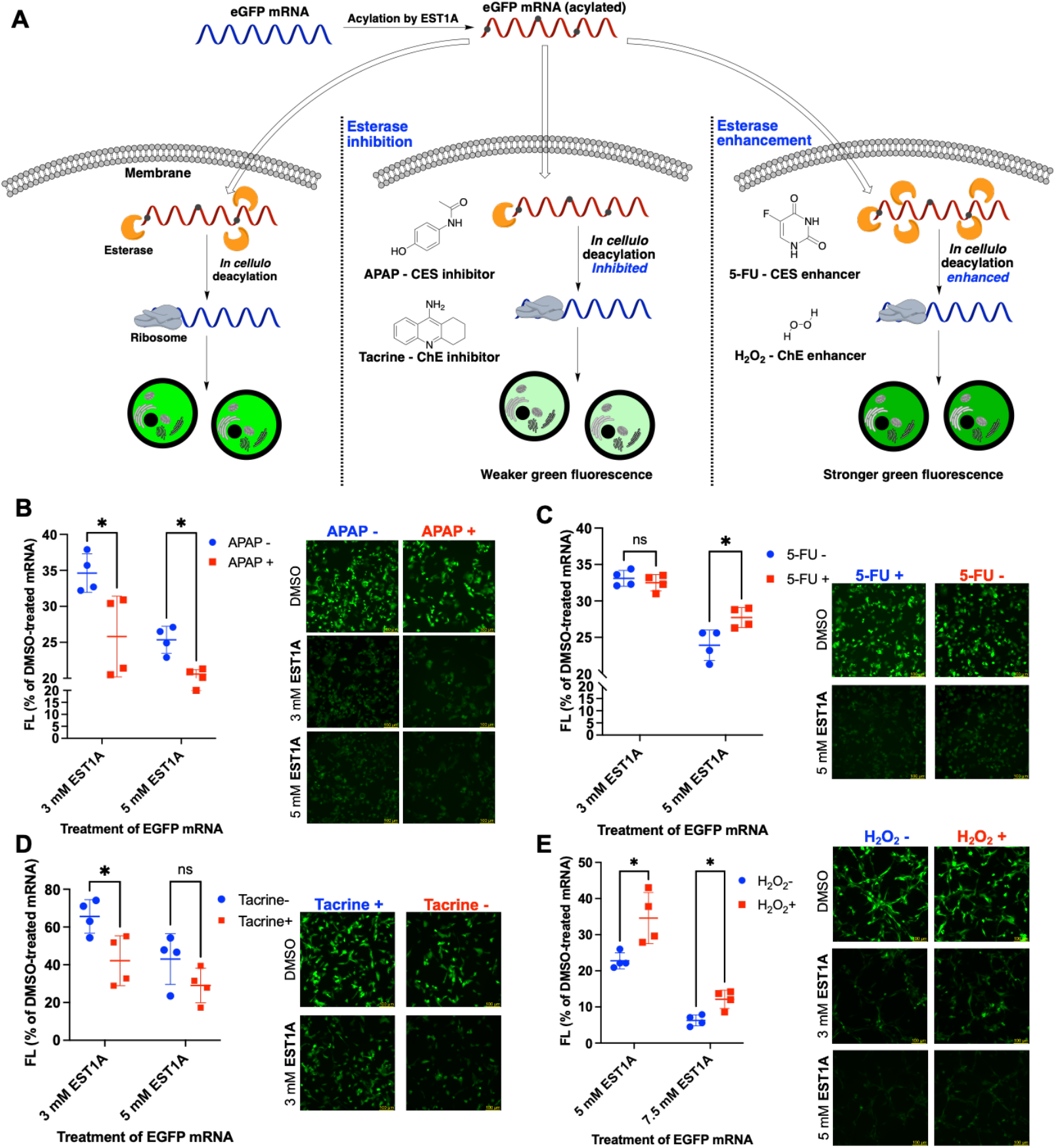
*In cellulo* deacylation of EGFP mRNA. (A) Schematic of modulating intracellular esterase activity with inhibitors and enhancers to control *in cellulo* deacylation of **EST1A**-acylated EGFP mRNA. (B, C) Quantification of EGFP fluorescence and representative HepG2 cell images for carboxylesterase inhibition (B) and enhancement (C). (D, E) Quantification of EGFP fluorescence and representative U87MG cell images for cholinesterase inhibition (D) and enhancement (E). Data are represented as mean ± SD (n = 4) with individual data points. Scale bars: 100 μm. t-test: ns, p > 0.05; *p ≤ 0.05; **p ≤ 0.01; ***p ≤ 0.001; ****p ≤ 0.0001. 40 ng/µL EGFP mRNA was acylated with 3, 5, or 7.5 mM **EST1A** in 10% DMSO at 37 °C for 4 h.

Guided by the inhibitory and enhancement assays, which established a positive correlation between intracellular CES activity and restoration of RNA function, we next investigated cell-specific deacylation in different cell lines. As a proof of concept, HEK293T cells were chosen as a noncancerous reference and compared with several cancer-derived lines (A549, MCF-7, U87MG, and HeLa). mCherry mRNA, encoding a red fluorescent protein, was used as a reporter and acylated with **EST1A** *in vitro* (Figure 6A). CES activity was measured using *p*-nitrophenyl acetate (*p*-NPA), whose hydrolyzed product absorbs at 405 nm. To enable comparison of cell specificity, lysates were prepared by lysing 1 million cells in 20 µL buffer for each cell line, and kinetic assays were then carried out with 250 µM p-NPA at a final lysate concentration of 50 cell-equivalents/µL. Kinetic analysis revealed the highest CES activity in U87MG cells, followed by HeLa, with MCF-7 and A549 exhibiting similar intermediate activity. All cancer cell lines showed stronger hydrolytic activity than HEK293T cells, consistent with the expected CES expression profiles of malignant versus noncancerous cells. The acylated mRNA was delivered into each line using Lipofectamine MessengerMAX. Cells were imaged the following day, and mCherry fluorescence was quantified; for normalization, the signal from cells transfected with DMSO-treated mRNA was defined as 100% in each cell line. These trends are summarized in the heat map in Figure 6B (also see Figure S8), which highlights a clear gradient of higher normalized mCherry fluorescence in the cancer cell lines relative to HEK293T across increasing **EST1A** concentrations. Moreover, the extent of RNA functional recovery closely followed the order of CES activity: U87MG and HeLa cells exhibited the greatest translational recovery, followed by MCF-7 and A549, with HEK293T showing the lowest response. Importantly, the acylated RNA caused only minimal cytotoxicity under the conditions tested (Figure S9). The difference between cancer and noncancerous cells indicates preferential restoration of reporter function in malignant cells, supporting the use of elevated CES expression as an endogenous trigger. These differences between malignant and noncancerous cells, across cancer lines derived from distinct tissues, indicate that **EST1A** can be leveraged for cancer-selective and tissue-specific activation of RNA-based therapeutics.

**Figure 6.**
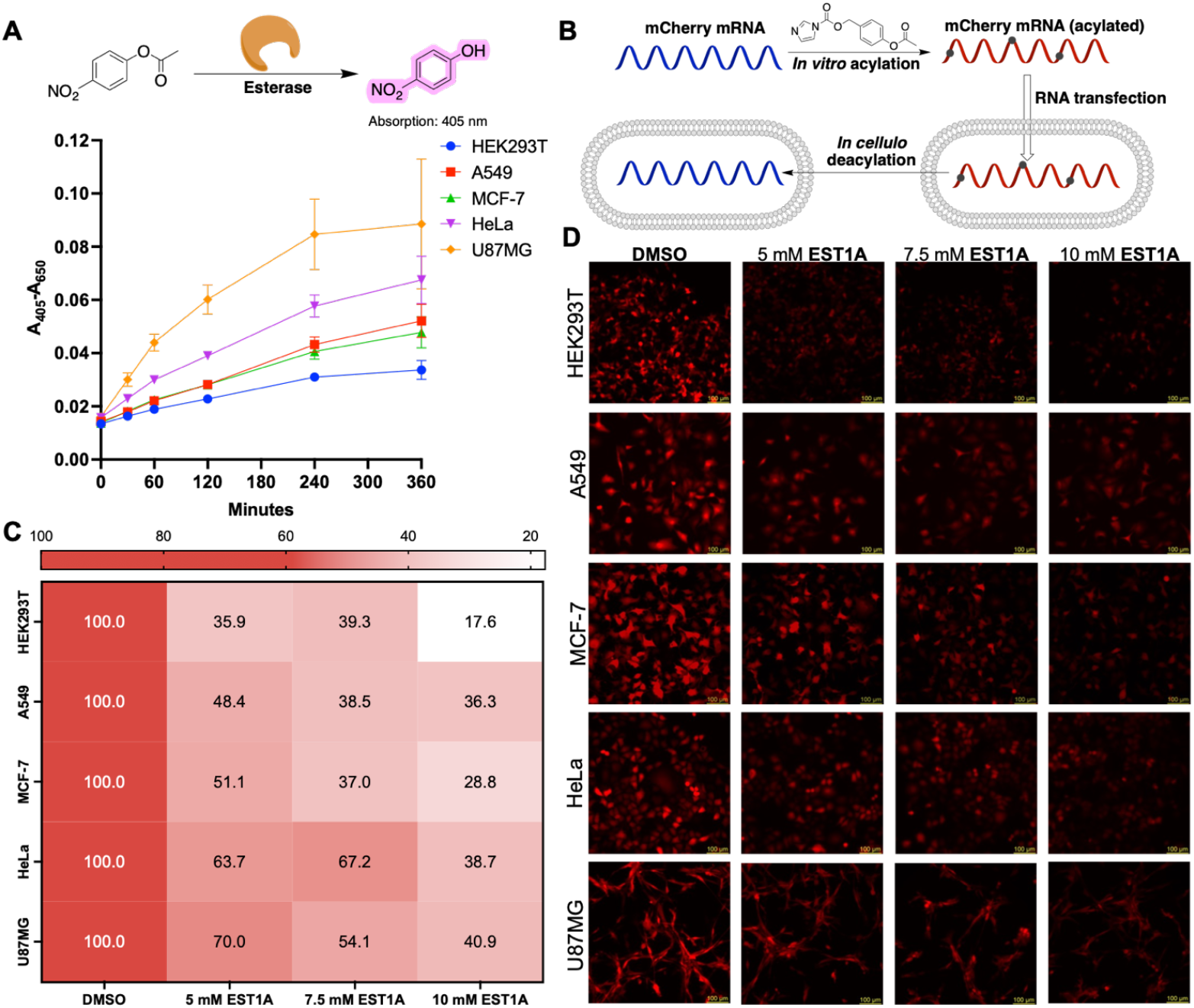
Specificity of *in cellulo* deacylation. (A) Kinetic analysis of p-NPA hydrolysis reporting esterase activity in lysates from different cell lines. Absorbance at 405 nm was monitored with 750 nm as the reference wavelength. Data are shown as mean ± SD (n = 4). (B) Schematic of *in cellulo* deacylation of **EST1A**-acylated mCherry mRNA. (C) Quantification of mCherry mRNA functional restoration; numbers indicate mean normalized fluorescence intensity (n = 4). (D) Representative fluorescence microscopy images of cell lines transfected with 0.3 ng/µL DMSO-treated or EST1A-acylated mCherry mRNA. Scale bar: 100 µm. For acylation, 40 ng/µL mCherry mRNA was incubated with 5, 7.5, or 10 mM **EST1A** in 10% DMSO at 37 °C for 2 h prior to transfection.

In summary, we have developed **EST1A** as a hydrolysis-responsive 2′-OH acylating reagent that enables reversible control of diverse RNAs, from short oligonucleotides to reporter mRNAs and aptamer. The RNA adducts are efficiently removed by endogenous esterases and histidine under benign conditions, avoiding the nonphysiological triggers commonly used in earlier reversible acylation systems. By combining modulation of carboxylesterase and cholinesterase activity with comparisons across noncancerous and cancer-derived cell lines, we uncover a direct link between intracellular esterase activity and functional recovery of acylated RNA, which allows **EST1A**-modified mRNA to function in a cell-selective and enzyme-dependent manner. We expect that hydrolysis-responsive 2′-OH acylation can be applied to build esterase-activated RNA therapeutics. Related designs that respond to other enzymes or metabolites could further extend this stimulus-responsive control of RNA.

## Supporting Information

Details of acylating reagent synthesis, RNA/oligonucleotide sequences, reversible acylation procedures, cell culture and assay conditions, additional gels and cell images, and spectroscopic data are provided in the Supporting Information.

## Acknowledgements

We thank the Department of Chemistry at the Mellon College of Science, Carnegie Mellon University (CMU), for support. This work was supported by the Startup Award (MCB200159) and used the Extreme Science and Engineering Discovery Environment (XSEDE), which is supported by the National Science Foundation (NSF) under grant no. ACI-1548562. We also gratefully acknowledge funding from NSF-MRI grant no. 2117784. We thank Xiaowei Ma (Collins lab, CMU) for valuable advice on the kinetic experiments and Mariah Arral (Whitehead lab, CMU) for sharing the A549 and HepG2 cell lines.

