## Supporting information for "RNA functional control by hydrolysis reversible acylation"

### 1. List of reagents and instruments

| **Chemical synthesis** | |
|---|---|
| 4-hydroxybenzyl alcohol | TCI |
| Acetic anhydride | Sigma-Aldrich |
| Triethylamine | Alfa Aesar |
| Imidazole | Sigma-Aldrich |
| chloroimidazole | Alfa Aesar |
| Sodium carbonate | Sigma-Aldrich |
| Sodium bicarbonate | Sigma-Aldrich |
| Sodium chloride | Fisher Scientific |
| Sodium sulfate | Fisher Scientific |
| Magnesium sulfate | Sigma-Aldrich |
| Potassium permanganate | Amazon |
| Triphosgene | Sigma-Aldrich |
| Dichloromethane (DCM) | Fisher Scientific |
| Hexane (Hex) | Fisher Scientific |
| Ethyl acetate (EtOAc) | Fisher Scientific |
| Methanol, HPLC grade | Fisher Scientific |
| Acetonitrile | Fisher Scientific |
| Deuterated chloroform ($CDCl_3$) | Sigma-Aldrich |
| Dimethyl sulfoxide (DMSO), anhydrous | Sigma-Aldrich |
| **Bioreagents** | |
| Water | Corning |
| Ethanol, 200 proof | Pharmco |
| Glycerol | Sigma-Aldrich |
| Orange G | Sigma-Aldrich |
| Bromophenol blue | Sigma-Aldrich |
| Xylene Cyanol FF | Sigma-Aldrich |
| Formamide | Alfa Aesar |
| Porcine liver esterase | Sigma-Aldrich |

| | |
|---|---|
| 1.0 M Tris-HCl buffer pH 7.5 | Sigma-Aldrich |
| 1.0 M Tris-HCl buffer pH 8.0 | Sigma-Aldrich |
| 1.0 M HEPES buffer | Gibco |
| 10 × PBS buffer pH 7.4 | Gibco |
| RNaseOUT™ Recombinant Ribonuclease Inhibitor | Invitrogen |
| Sodium acetate buffer solution, pH 5.2±0.1 (25 °C), for molecular biology, 3 M | Sigma-Aldrich |
| Glycogen, 5 mg/mL | Invitrogen |
| SYBR-Gold | Invitrogen |
| TEMED | National diagnostic |
| SequaGel – UreaGel Buffer | National diagnostic |
| SequaGel – UreaGel Diluent | National diagnostic |
| SequaGel UreaGel 19:1 Concentrate | National diagnostic |
| Urea | Acros Organics |
| Ammonium persulfate | Sigma-Aldrich |
| Agarose | Sigma-Aldrich |
| Acrylamide: Bis-Acrylamide 19:1 (40% Solution/Electrophoresis) | Fisher Scientific |
| Tris-Borate-EDTA, 10X Solution, Electrophoresis | Fisher Scientific |
| Cyanine 3 Phosphoramidite | Glen research |
| 2'-tBDSilyl Uridine CED phosphoramidite | Chemgene |
| 2'-tBDSilyl Uridine 3'-lcaa CPG 500Å | Chemgene |
| $MgCl_2$, 1 M | Invitrogen |
| KCl, 2 M | Invitrogen |
| Phusion High-Fidelity PCR Kit | Thermo Scientific |
| GeneJet Gel Extraction Kit | Thermo Scientific |
| MEGAshortscript™ T7 Transcription Kit | Invitrogen |
| CleanCap® EGFP mRNA | TriLink |

| | |
|---|---|
| CleanCap® mCherry mRNA | TriLink |
| Oligo Clean & Concentrator | Zymo research |
| Quick-RNA Miniprep Kit | Zymo research |
| Pierce™ BCA Protein Assay Kits | Thermo Scientific |
| **Cell culture & assays** | |
| Dulbecco's Modified Eagle Medium (DMEM) | Gibco |
| Eagle's Minimum Essential Medium (EMEM) | Corning |
| Opti-MEM™ I Reduced Serum Medium, no phenol red | Gibco |
| Insulin, human recombinant, zinc solution | Gibco |
| Lipofectamine™ 3000 | Invitrogen |
| Lipofectamine MessengerMAX | Invitrogen |
| 1 × PBS buffer pH 7.4 | Gibco |
| 0.25% Trypsin-EDTA (1 ×) | Gibco |
| Fetal bovine serum, premium, USA origin | Corning |
| Penicillin-Streptomycin (10,000 U/mL) | Gibco |
| Sodium pyruvate | Gibco |
| Trypan blue stain | Gibco |
| Acetaminophen | Johnson & Johnson |
| 5-fuorouracil | Thermo Scientific |
| DMSO | ATCC |
| Tacrine | Sigma-Aldrich |
| Hydrogen peroxide | Sigma-Aldrich |
| *p*-Nitrophenyl Acetate | TCI |
| SuperScript™ III First-Strand Synthesis System | Invitrogen™ |
| PowerTrack™ SYBR Green Master Mix for qPCR | Applied Biosystems™ |
| LIVE/DEAD™ Cell Vitality Assay Kit, C12 Resazurin/SYTOX™ Green | Invitrogen |

| **Instruments** | |
|---|---|
| Nuclear Magnetic Resonance Spectrometer | Bruker Avance™ III 500 MHz and Bruker NEO 500 MHz NMR |
| Mass spectrometer | Thermo Scientific™ Exactive™ Plus EMR mass spectrometer |
| Gel imager | BIO-RAD ChemiDoc MP Imaging System |
| Quantitative PCR | QuantStudio™ 5 Real-Time PCR System |
| Confocal microscope | Leica DMi8 |
| DNA/RNA Synthesizer | Mermade 4 |
| Plate reader | Tecan Spark |
| MALDI TOF | Bruker Ultraflextreme MALDI TOF/TOF MS |

### 2. Synthesis and molecular characterization

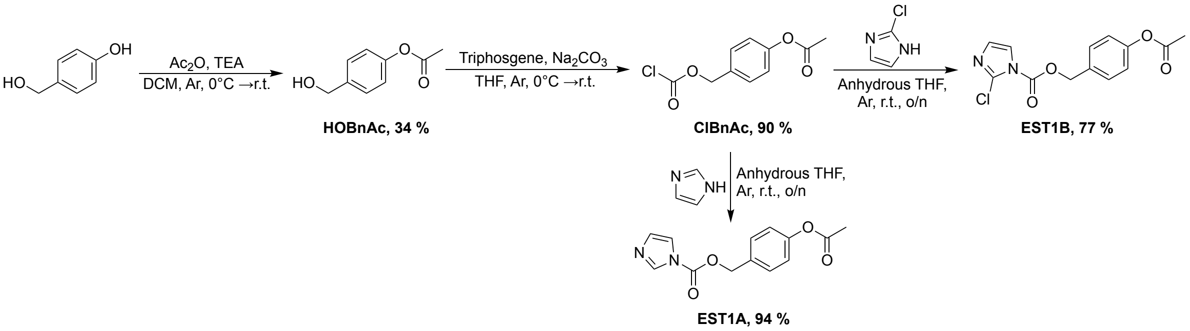


**Scheme S1.** Synthetic routes of acylating reagents.

*4-(hydroxymethyl)phenyl acetate* (**HOBnAc**)

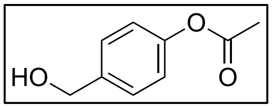

To a solution of 497 mg (4.0 mmol, 1.0 eq) 4-hydroxybenzyl alcohol and 669 µL (4.8 mmol, 1.2 eq) triethylamine in 4 mL DCM under Ar was added a solution of 454 µL (4.8 mmol, 1.2 eq) $Ac_2O$ in 4 mL DCM dropwisely at 0°C. The reaction was next warmed to room temperature, stirred overnight, and concentrated under vacuo. The residue was redissolved in EtOAc and washed with $NaHCO_3$ and brine. The organic layer was dried over $MgSO_4$, filtered, and concentrated under vacuo. The crude product was finally purified by flash column chromatography (Hex/EtOAc = 3:1, v/v). The light-yellow oil was yielded. 225 mg, 34%. The synthesis was repeated to obtain enough product for later steps. $^1$H NMR (500 MHz, $CDCl_3$) δ 7.41 – 7.35 (m, 2H), 7.12 – 7.05 (m, 2H), 4.68 (s, 2H), 2.31 (s, 3H). $^{13}$C NMR (126 MHz, $CDCl_3$) δ 169.58, 149.20, 140.33, 128.62, 121.66, 65.60, 22.43.

*4-(acetoxy)benzyl chloroformate* **(ClBnAc)**

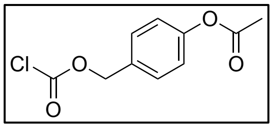

To a solution suspension of 538 mg (5.08 mmol, 4.0 eq) sodium carbonate and 377 mg (1.27 mmol, 1.0 eq) triphosgene in 6 mL anhydrous THF under Ar in the ice bath, was added a solution of 211 mg (1.27 mmol, 1.0 eq) **HOBnAc** in 6 mL anhydrous THF dropwisely. After the addition, the reaction was warmed to room temperature and stirred overnight. THF was then evaporated under vacuo. The residue was redissolved in Hex/EtOAc = 9:1 (v/v) and filtered by a 2 cm silica gel pad. The filtrate was concentrated under vacuo to yield yellow oil. 178 mg, 61%. The synthesis was repeated to obtain enough product for the next step. $^1$H NMR (500 MHz, $CDCl_3$) δ 7.43 (d, $J$ = 8.6 Hz, 2H), 7.15 (d,

$J$ = 8.5 Hz, 2H), 5.30 (s, 2H), 2.33 (s, 3H). $^{13}$C NMR (126 MHz, $CDCl_3$) δ 169.68, 151.39, 150.59, 130.86, 130.25, 122.06, 72.62, 21.50.

*4-acetoxybenzyl 1H-imidazole-1-carboxylate* **(EST1A)**

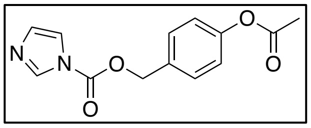

Under Ar, to a solution of 30 mg (0.13 mmol, 1.0 eq) **ClBnAc** in 1 mL anhydrous THF, was added a solution of 18 mg (0.26 mmol, 2.0 eq) imidazole in 1 mL anhydrous THF dropwisely. The reaction was stirred at room temperature overnight before filtration. The filtrate was then concentrated under vacuum to yield yellow oil. 32 mg, 94 %. The synthesis was repeated to obtain enough product for the acylation study. $^{1}$H NMR (500 MHz, $CDCl_3$) δ 8.16 (s, 1H), 7.48 (d, $J$ = 8.5 Hz, 2H), 7.44 (s, 1H), 7.16 (d, $J$ = 8.5 Hz, 2H), 7.08 (s, 1H), 5.42 (s, 2H), 2.33 (s, 3H). $^{13}$C NMR (126 MHz, $CDCl_3$) δ 169.22, 151.72, 148.50, 137.15, 131.52, 130.76, 130.09, 122.10, 117.13, 69.52, 21.07. ESI-MS: $[M+H]^+$ calculated $m/z$ 261.0876; found 261.0863.

*4-acetoxybenzyl 2-chloro-1H-imidazole-1-carboxylate* **(EST1B)**

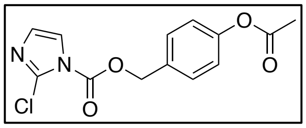

Under Ar, to a solution of 37 mg (0.16 mmol, 1.0 eq) **ClBnAc** in anhydrous THF, was added a solution of 33 mg (0.32 mmol, 2.0 eq) chloroimidazole in anhydrous THF dropwisely. The reaction was stirred at room temperature overnight before filtration. The filtrate was then concentrated under vacuum to yield yellow oil. 37 mg, 77%. The synthesis was repeated to obtain enough product for the acylation study. $^{1}$H NMR (500 MHz, $CDCl_3$) δ 7.49 (d, $J$ = 8.5 Hz, 2H), 7.46 (d, $J$ = 1.8 Hz, 1H), 7.16 (d, $J$ = 8.5 Hz, 2H), 6.94 (d, $J$ = 1.8 Hz, 1H), 5.42 (s, 2H), 2.33 (s, 3H). $^{13}$C NMR (126 MHz, $CDCl_3$) δ 169.24, 151.27, 147.75, 133.06, 131.26, 130.16, 128.72, 122.13, 120.23, 69.54, 21.10. ESI-MS: $[M+K]^+$ calculated $m/z$: 333.0045; found 333.0036.

### 3. Procedure for reversible acylation

**DNA and RNA sequences.**

**Cy3-$U_{16}$ oligonucleotide:** 5′-Cy3-rUrUrU rUrUrU rUrUrU rUrUrU rUrUrU rU-3′, synthesized by DNA/RNA synthesizer.

**EGFP mRNA:** CleanCap® EGFP mRNA (5moU)- (L-7201) from TriLink

**mCherry mRNA**: CleanCap® mCherry mRNA (5moU) - (L-7203) from TriLink

**Antisense strand of EGFP siRNA:** customize synthesis by IDT, 5′-rUrCrC rUrUrG rArArG rArArG rArArG rArUrG rGrUrG rCdTdT-3′.

**Scrambled sequence:** customize synthesis by IDT, 5′-rGrGrU rGrCrG rArArU rUrGrG rUrCrA rCrUrA rAdTdT-3′.

**F-30 broccoli aptamer DNA template:**

5′_TAATACGACTCACTATAGGGTTGCCATGTGTATGTGGGAGACGGTCGGGTCCAGA
TATTCGTATCTGTCGAGTAGAGTGTGGGCTCCCACATACTCTGATGATCCTTCGGGA
TCATTCATGGC_3′

**Primers of F-30 broccoli aptamer:**

5′-TAATACGACTCACTATAGGGTTGC-3′ (forward)

5′-GCCATGAATGATCCCGAAG-3′ (reverse)

**Primers of cDNA of EGFP mRNA:**

5′-TCAAGTCCGCCATGCCC-3′ (forward)

5′-CCGCCCTATAATGGGCGGAGGAG-3′ (reverse)

**Primers of GADPH:**

5′-ACCACAGTCCATGCCATCAC-3′ (forward)

5′-TCCACCACCCGTTTGCTGTA-3′ (reverse)

**Ethanol precipitation.** After the incubation, the reaction mixture was brought to 86.5 μL with water, then 9.9 μL sodium acetate (3 M, pH 5.2), 4.6 μL glycogen (5 mg/mL), and 404 μL ethanol were added. The mixture was kept at -80 °C overnight before centrifugation at 4 °C (30 min, 20600 rcf). The supernatant was removed by aspiration, and the pellet was air-dried for 10 min before resuspension.

**2′-OH acylation of Cy3-$U_{16}$ RNA.** The reaction volume is 10 μL. In a centrifuge tube, 0.65 ng/μL (120 nM) Cy3-$U_{16}$ RNA was incubated with **EST1A** (50-100 mM), **EST1B** (100 mM) in 10%-40%DMSO (conditions shown in the figure legends). After incubation, the reaction was mixed with 10 μL orange G in formamide (loading dye). The sample was denatured by heating the RNA at 90 °C for 3 min and immediately cooling it in an ice bath. Finally, the reaction was analyzed by 30% denaturing polyacrylamide gel electrophoresis (PAGE, monomer ratio: 19:1). The gel was visualized by the Cy3 fluorescence channel.

**Reversible acylation of Cy3-$U_{16}$ RNA.** For acylation, 0.65 ng/μL (120 nM) Cy3-$U_{16}$ oligonucleotide was incubated with 50 mM **EST1A** in 10% DMSO at 37 °C for 6 hours (reaction volume: 10 μL) and then precipitated by EtOH. For deacylation by histidine: the RNA pellet was resuspended in 50 mM histidine in 1 × PBS buffer (pH 7.4). For deacylation by imidazole the RNA pellet was resuspended in 50 mM imidazole in 50 mM HEPES buffer (pH 7.5). For deacylation by porcine liver esterase: the RNA pellet was resuspended in 40 U/mL in 50 mM Tris-HCl buffer (pH 8.0). The volume of deacylation is 10 μL with 120 nM RNA concentration. After incubating at 37 °C for 4 hours, the reaction was mixed with 10 μL orange G in formamide, denatured, and analyzed by 30% denaturing PAGE as previously described.

**Reversible acylation of Antisense strand of EGFP siRNA (siEGFP-AS).** For acylation: 5 μM (100 pmol in 20 μL) siEGFP-AS incubated with 50 mM **EST1A** in 10% DMSO at 37 °C for 4 hours and then precipitated by EtOH. For deacylation, 5 μM (100 pmol in 20 μL) acylated siEGFP-AS incubated with 40 U/mL porcine liver esterase (PLE) or HeLa cell lysate (total protein: 0.5 μg/μL) in 50 mM Tris-HCl (pH 8.0) at 37 °C for 4 hours. The reaction was mixed with 20 μL orange G in formamide, denatured, and analyzed by 30% denaturing PAGE as previously described.

**MALDI TOF analysis of siEGFP-AS.** For acylation, 5 μM siEGFP-AS incubated with 20 mM EST1A in 10% DMSO at 37 for 4 hours and precipitated by EtOH. For deacylation, 5 μM acylated siEGFP-AS incubated with 40 U/mL porcine liver esterase (PLE) in 50 mM Tris-HCl (pH 8.0) at 37 °C for 4 hours and precipitated by EtOH. The RNA pellet needs to be washed with 500 μL 75% EtOH twice and 500 μL 100% EtOH once to remove the salt. The matrix was prepared by mixing 9 volumes of saturated 3-hydroxypicolinic acid in TA50 (water/acetonitrile = 1:1 with 0.1% TFA) with 1 volume of 100 mg/mL ammonium citrate dibasic in water. Add 1.5 μL of matrix to the plate and let it dry in the air. Then, 2 μL of RNA in water was added to the matrix crystal and allowed to dry in the air. The addition of RNA was repeated until the desired amount of RNA was added to the plate. We added 100 pmol RNA in our experiments.

**Production of F-30 broccoli aptamer.** The DNA template of the aptamer (100 ng) was amplified by Phusion High-Fidelity PCR Kit (conditions shown in the table below), purified by 2% agarose gel, and then extracted by GeneJet Gel Extraction Kit. The aptamer was then produced by *in vitro* transcription with 500 ng of template using MEGAshortscript™ T7 Transcription Kit and isolated by EtOH precipitation. The length and purity of aptamer were analyzed with 2% agarose gel.

| | | |
|---|---|---|
| 98 ˚C | | 30 s |
| 30 Repeats | 98 ˚C | 10 s |
| | 60 ˚C | 30 s |
| | 72 ˚C | 15 s |
| 72 ˚C | | 5 min |
| 4 ˚C | | Forever |

**Reversible acylation of F-30 broccoli aptamer.** For acylation, 5 μg F-30 broccoli aptamer was incubated with **EST1A** in 10% DMSO (or only DMSO) at 37 °C for 4 hours (reaction volume: 25 μL) and purified by Oligo Clean & Concentrator. For deacylation by histidine, 5 μg of F-30 broccoli aptamer was incubated with 50 mM histidine in 1 × PBS (pH 7.4) at 37 °C for 4 hours (reaction volume: 50 μL) and purified by Oligo Clean & Concentrator. For deacylation by imidazole, broccoli aptamer was incubated with 50 mM HEPES buffer (pH 7.5) at 37°C for 4 hours. RNAs after deacylation were purified by Oligo Clean & Concentrator.

**Aptamer-DFHBI fluorescence assay.** 5 μg acylated or DMSO-treated aptamer in 250 μL was first folded in the HEPES folding buffer (40 mM HEPES, 100 mM KCl, 5 mM $MgCl_2$, pH = 7.5) by heating in the heat block to 90 °C for 3 min and then placing the heat block on the bench until the temperature dropped to room temperature. Then 2.5 μL of 1 mM DFHBI (in DMSO/$H_2O$ = 1:19) was added to the folded aptamer, and the mixture was next incubated at 37°C for 1 h. For quantitative analysis, the fluorescence intensity was measured by the plate reader (excitation: 469 nm, width 5 nm; emission: 501 nm, width 5 nm).

**Reversible acylation of EGFP mRNA.** For acylation, 40 ng/µL EGFP mRNA was incubated with **EST1A** in 10% DMSO at 37 °C for 4 hours (reaction volume: 10 µL) and precipitated by EtOH. The DMSO-treated or acylated RNA was resuspended to the desired concentration to study deacylation *in cellulo*. For deacylation by histidine, the 200 ng of mRNA pellet was incubated with 50 mM histidine in 1 × PBS (pH 7.4) at 37 °C for 4 hours (reaction volume: 10 µL). For deacylation by imidazole, the mRNA pellet was incubated with 50 mM imidazole in 50 mM HEPES (pH 7.5) at 37°C for 4 hours. The mRNA was collected by EtOH precipitation and then resuspended to 20 ng/µL for transfection.

**Cell culture.** HeLa and HEK293T cells were cultured with DMEM culture medium (10% FBS, 1% Penicillin/Streptomycin). HepG2 cells were cultured with DMEM culture medium (15% FBS, 1% Penicillin/Streptomycin). U87MG cells were cultured with EMEM culture medium (10% FBS, 1% Penicillin/Streptomycin). MCF-7 cells were cultured with EMEM culture medium (10% FBS, 1% Penicillin/Streptomycin, 10 µg/mL insulin). Both cells are incubated at 37°C and 5% $CO_2$ with 95% humidity. For subculture, the old medium was removed when cells reached 90% confluency. After washing with PBS, cells were dissociated by trypsin-EDTA (0.25%). Trypsinization was quenched by adding fresh culture medium, and a quarter of the cells were transferred to a new vessel.

**Transfection of EGFP mRNA for the study of *in vitro* reversible acylation.** 20 k/well HepG2 cells were seeded in a 96-well plate with DMEM culture medium (15% FBS, 1% Penicillin/Streptomycin) and incubated overnight at 37°C and 5% $CO_2$ with 95% humidity. Before the transfection, the old media was replaced with 100 µL fresh DMEM medium. For each well, 5 µL OptiMEM medium was mixed with 0.2 µL Lipofectamine MessengerMAX and incubated for 15 min at room temperature. 50 ng EGFP mRNA in 2.5 µL RNase-free water was diluted with 5 µL OptiMEM medium and incubated at room temperature for 5 min. The mRNA/OptiMEM and Lipofectamine/OptiMEM mixtures were then combined and incubated for 10 min at room temperature. The final mixture was dropwisely added into each well. 4 hours later, the medium was replaced with 100 µL fresh DMEM medium. After incubating overnight, the cells were imaged in phenol red-free DMEM culture medium (15% FBS, 1% Penicillin/Streptomycin) by the phase and GFP channels of the Leica DMi8 confocal microscope (GFP excitation wavelength: 450-490 nm, emission wavelength: 500-550 nm). For quantitative analysis, the fluorescence intensity was measured by plate reader (excitation: 488 nm, width 5 nm; emission: 509 nm, width 5 nm). Cells without mRNA transfection were measured as blank.

**Co-transfection of EGFP-AS and EGFP mRNA.** 40 k/well HepG2 cells were seeded in a 48-well plate with DMEM culture medium (15% FBS, 1% Penicillin/Streptomycin) and incubated overnight at 37°C and 5% $CO_2$ with 95% humidity. Before the transfection, the old media was replaced with 180 µL fresh DMEM medium. For each well, 10 µL OptiMEM medium was mixed with 0.4 µL Lipofectamine 3000; 66.7 ng EGFP mRNA (0.667 µL) and 20 pmol siRNA (4 µL) were mixed, diluted with 10 µL OptiMEM medium, and incubated at room temperature for 5 min. The RNA/OptiMEM and Lipofectamine/OptiMEM mixtures were then combined and incubated for 15 min at room temperature. The final mixture was dropwisely added into each well. 4 hours later, the medium was replaced with 200 µL fresh medium. After incubating overnight, the cells were imaged by the phase and GFP channels of the Leica DMi8 confocal microscope (GFP excitation wavelength: 450-490 nm, emission wavelength: 500-550 nm). For quantitative analysis, the fluorescence intensity in each image was calculated by CellProfiler. Experiments were performed in quadruplicate (biological repeats), and six images (technical repeats) were taken in each well.

**RT-qPCR analysis of EGFP mRNA degradation.** Followed by the fluorescent image in 48-well plate, total RNA was extracted from cells using the Quick-RNA Miniprep Kit according to the manufacturer's protocol. From each sample, 400 ng of RNA was reverse-transcribed to synthesize cDNA using the SuperScript™ III First-Strand Synthesis System in a 20 µL reaction volume. The resulting cDNA was diluted 100-fold, and 1 µL was used as a template for qPCR of 10 µL. The reactions were performed with PowerTrack™ SYBR Green Master Mix (conditions shown in the table below). The analysis quantified the expression of EGFP relative to the endogenous housekeeping gene, GADPH. All experiments were conducted with four biological replicates, and each qPCR reaction was run with eight technical replicates (10 µL each).

| | |
|---|---|
| 95 ˚C | 10 min |

<table>
<tr><td rowspan="2">40 Repeats</td><td>95 ˚C</td><td>15 s</td></tr>
<tr><td>60 ˚C</td><td>60 s</td></tr>
<tr><td colspan="2">60 ˚C</td><td>60 s</td></tr>
<tr><td colspan="2">95 ˚C</td><td>15 s</td></tr>
<tr><td colspan="2">4 ˚C</td><td>Forever</td></tr>
</table>

***In vitro* acylation of EGFP mRNA.** 40 ng/µL EGFP mRNA was incubated with 3, 5 or 7.5 mM **EST1A** in 10% DMSO at 37 °C for 4 hours (reaction volume: 10 µL) and precipitated by EtOH. The mRNA was collected by EtOH precipitation and then resuspended to 16.7 ng/µL for transfection.

***In vitro* acylation of mCherry mRNA.** 40 ng/µL mCherry mRNA was incubated with 5, 7.5, 10 mM **EST1A** in 10% DMSO at 37 °C for 2 hours (reaction volume: 10 µL) and precipitated by EtOH. The mRNA was collected by EtOH precipitation and then resuspended to 16.7 ng/µL for transfection.

**Acetaminophen (APAP) and 5-fluorouracil treatment of HepG2 cells and *in cellulo* deacylation of EGFP mRNA.** 16 k/well HepG2 cells were seeded in a 96-well plate with DMEM culture medium (15% FBS, 1% Penicillin/Streptomycin) and incubated overnight at 37°C and 5% $CO_2$ with 95% humidity. The old medium was aspirated. For APAP, fresh medium containing 1 mM APAP (1 M stock in DMSO) or 0.1% DMSO was added. For 5-FU, fresh medium containing 80 µM 5-FU (100 mM stock in DMSO) or 0.08% DMSO was added. After 24 h, the old media was replaced with 100 µL fresh DMEM medium. For each well, 3.75 µL OptiMEM medium was mixed with 0.15 µL Lipofectamine MessengerMAX and incubated for 15 min at room temperature. 25 ng EGFP mRNA in 1.5 µL water was diluted with 5 µL OptiMEM medium and incubated at room temperature for 5 min. The mRNA/OptiMEM and Lipofectamine/OptiMEM mixtures were then combined and incubated for 10 min at room temperature. The final mixture was dropwisely added into each well (4 wells in total). 4 hours later, the medium was replaced with 100 µL fresh DMEM medium. After incubating overnight, the cells were imaged in phenol red-free DMEM culture medium (15% FBS, 1% Penicillin/Streptomycin) by the phase and GFP channels of the Leica DMi8 confocal microscope (GFP excitation wavelength: 450-490 nm, emission wavelength: 500-550 nm). For quantitative analysis, the area that cells occupy in each image was calculated by the software ilastik (pixel classification) and CellProfiler. The fluorescence intensity in each image was calculated by CellProfiler. The ratio of fluorescence intensity to the cell-occupying area was plotted (blank deducted). Experiments were performed in quadruplicate (biological repeats), and six images (technical repeats) were taken in each well.

**Tacrine and hydrogen peroxide ($H_2O_2$) treatment of U87MG cells and *in cellulo* deacylation of EGFP mRNA.** 16 k/well U87MG cells were seeded in a 96-well plate with EMEM culture medium (10% FBS, 1% Penicillin/Streptomycin) and incubated overnight at 37°C and 5% $CO_2$ with 95% humidity. The old medium was aspirated. For the inhibitory assay, fresh medium with and without 100 µM tacrine (50 mM stock in water) was added 2 hours before transfection. For the enhancive assay, fresh medium with and without 200 µM $H_2O_2$ (100 mM stock in water) was added 1 hour before transfection. The old medium was then replaced by 100 µL fresh medium right before the transfection (for the inhibitory assay, the fresh medium with 50 µM was added). For each well, 3.75 µL OptiMEM medium was mixed with 0.15 µL Lipofectamine MessengerMAX and incubated for 15 min at room temperature. 25 ng EGFP mRNA in 1.5 µL water was diluted with 5 µL OptiMEM medium and incubated at room temperature for 5 min. The mRNA/OptiMEM and Lipofectamine/OptiMEM mixtures were then combined and incubated for 10 min at room temperature. The final mixture was dropwisely added into each well (4 wells in total). 4 hours later, the medium was replaced with 100 µL fresh EMEM medium (for the inhibitory assay, fresh medium with 100 µM tacrine was added). After incubating overnight, the cells were imaged in phenol red-free DMEM culture medium (10% FBS, 1% Penicillin/Streptomycin) by the phase and GFP channels of the Leica DMi8 confocal microscope (GFP excitation wavelength: 450-490 nm, emission wavelength: 500-550 nm). For quantitative analysis, the area that cells occupy in each image was calculated by the software ilastik (pixel classification) and CellProfiler. The fluorescence intensity in each image was calculated by CellProfiler. The ratio of fluorescence intensity to the cell-occupying area was plotted (blank deducted). Experiments were performed in quadruplicate (biological repeats), and six images (technical repeats) were taken in each well.

**Preparation of cell lysate (cytoplasmic fraction).** Cell lysates were prepared following a previously reported procedure with modifications.[1] Cells were cultured as mentioned before. The growth medium was removed from the cells, and the cells were washed twice with 1× PBS. Cells were harvested by scraping in fresh PBS and pelleted by centrifugation at 450 × g for 5 minutes. The supernatant was discarded, and the cell pellet was resuspended in 5 × packed cell volume

(PCV) of lysis buffer (10 mM HEPES, 1.5 mM $MgCl_2$, 10 mM KCl) freshly supplemented with protease inhibitor cocktail tablet (one tablet for 40 mL buffer). After a 15-minute incubation on ice to allow for cell swelling, the suspension was centrifuged at 400 × g for 5 minutes at 4 °C, and the supernatant was discarded. The pellet was then resuspended in lysis buffer (1 million/20µL) and mechanically lysed by passing the suspension 14 times through a 25-gauge needle. The cells were centrifuged at 11,000 × g for 20 minutes at 4 °C. The supernatant was transferred to a fresh tube, and the protein concentration was determined by Pierce™ BCA Protein Assay Kits.

**Measurement of esterase level across cell lines.** For a 200 µL reaction, 188 µL of water, 10 µL of 1 M HEPES buffer (pH 7.5), and 1 µL of cell lysate (50,000 cell equivalent) were mixed, followed by the addition of 1 µL of 50 mM *p*-NPA. The reaction mixture was dispensed into a 96-well plate in quadruplicate (50 µL per well) and incubated at 37 °C. Absorbance at 405 nm was monitored with a reference wavelength of 650 nm.

***In cellulo* deacylation of mCherry mRNA in different cell lines.** 20 k/well cells were seeded in a 96-well plate with corresponding culture medium and incubated overnight at 37°C and 5% $CO_2$ with 95% humidity. The old medium was aspirated and replaced by 100 µL fresh medium right before the transfection. For each well, 3.75 µL OptiMEM medium was mixed with 0.15 µL Lipofectamine MessengerMAX and incubated for 15 min at room temperature. 25 ng mCherry mRNA in 1.5 µL water was diluted with 5 µL OptiMEM medium and incubated at room temperature for 5 min. The mRNA/OptiMEM and Lipofectamine/OptiMEM mixtures were then combined and incubated for 10 min at room temperature. The final mixture was dropwisely added into each well (4 wells in total). 4 hours later, the medium was replaced with 100 µL fresh medium. After incubating overnight, the cells were imaged by the phase and GFP channels of the Leica DMi8 confocal microscope (GFP excitation wavelength: 450-490 nm, emission wavelength: 500-550 nm). For quantitative analysis, the area that cells occupy in each image was calculated by the software ilastik (pixel classification) and CellProfiler. The fluorescence intensity in each image was calculated by CellProfiler. The ratio of fluorescence intensity to the cell-occupying area was plotted (blank deducted). Experiments were performed in quadruplicate (biological repeats), and six images (technical repeats) were taken in each well.

**Cytotoxicity test.** 20 k/well U87MG or HepG2 cells were seeded in a 96-well plate with the corresponding culture medium and incubated overnight at 37°C and 5% $CO_2$ with 95% humidity. The old medium was aspirated. For each well, 5 μL OptiMEM medium was mixed with 0.2 μL Lipofectamine MessengerMAX and incubated for 15 min at room temperature. 10 pmol siRNA (2 μL) was diluted with 5 μL OptiMEM medium and incubated at room temperature for 5 min. The siRNA/OptiMEM and Lipofectamine/OptiMEM mixtures were then combined and incubated for 10 min at room temperature. The final mixture was dropwisely added into each well. 4 hours later, the old was removed by aspiration. The cell viability was tested by LIVE/DEAD™ Cell Vitality Assay Kit following the manufacturer's protocol.

4. **Additional gel images, cell images, and spectra.**

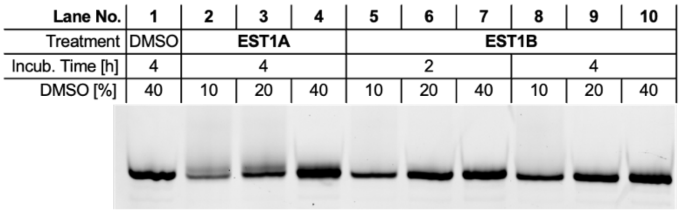


Figure S1. Acylation of 120 nM Cy3-$U_{16}$ oligonucleotide by 100 mM **EST1A** and **EST1B** in 10-40% DMSO at 25 °C for 2 or 4 hours.

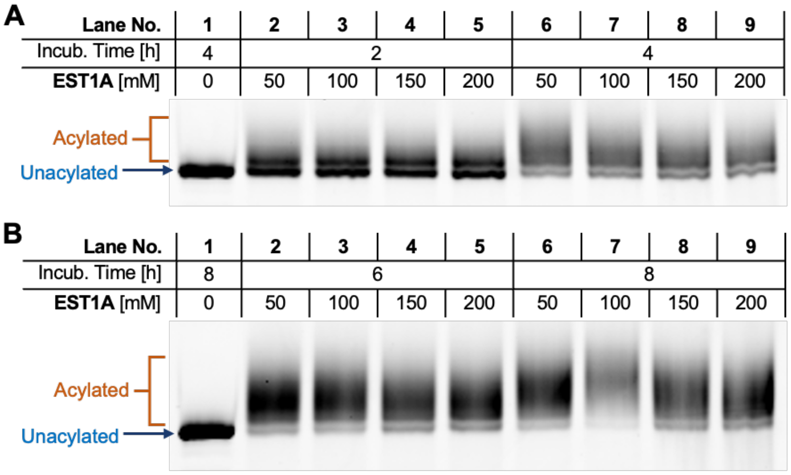


Figure S2. Acylation of 120 nM Cy3-$U_{16}$ oligonucleotide by 50-200 mM **EST1A** in 10% DMSO at 37 °C for 2-8 hours.

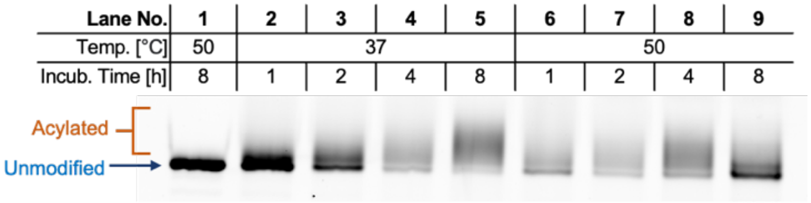


Figure S3. Acylation of 120 nM Cy3-$U_{16}$ oligonucleotide by 100 mM **EST1A** in 10% DMSO at 37 and 50 °C for 1-8 hours.

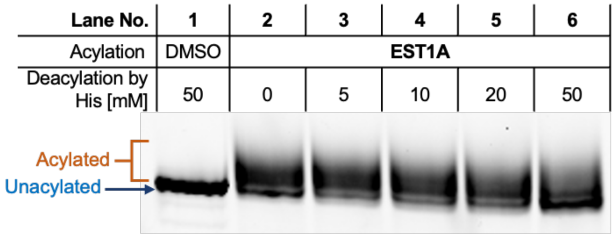


Figure S4. Reversible acylation of Cy3-$U_{16}$ oligonucleotide. Acylation: 120 nM RNA oligonucleotide and 50 mM **EST1A** incubated in 10% DMSO at 37°C for 6 h. Deacylation: 120 nM acylated RNA oligonucleotide incubated with 50 mM histidine in 1 × PBS (pH 7.4) at 37°C for 4 h.

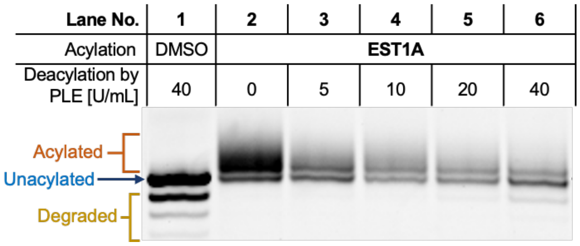


Figure S5. Reversible acylation of Cy3-$U_{16}$ oligonucleotide. Acylation: 120 nM RNA oligonucleotide and 50 mM **EST1A** incubated in 10% DMSO at 37°C for 6 h. Deacylation: 120 nM acylated Cy3-$U_{16}$ RNA oligonucleotide incubated with porcine liver esterase in 50 mM Tris-HCl buffer (pH 8.0) at 37°C for 4 h

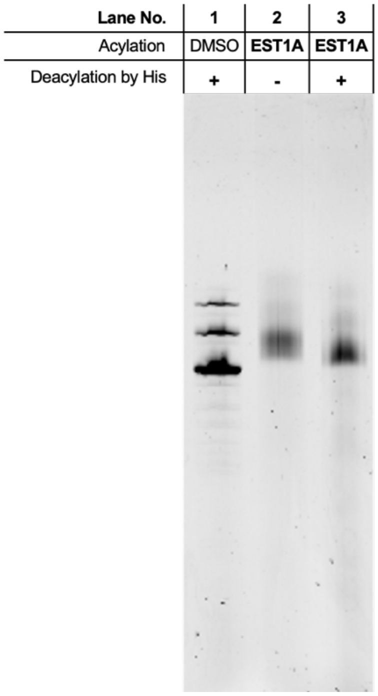


Figure S6. Reversible acylation of broccoli aptamer analyzed by 12% dPAGE. Acylation: 100 ng/µL broccoli aptamer and 50 mM **EST1A** incubated in 10% DMSO at 37°C for 4 h. Deacylation: 100 ng/µL acylated broccoli aptamer and 50 mM histidine incubated in 1 × PBS buffer (pH 7.4) at 37°C for 4 h. The gel was scanned in the SYBR-gold channel.

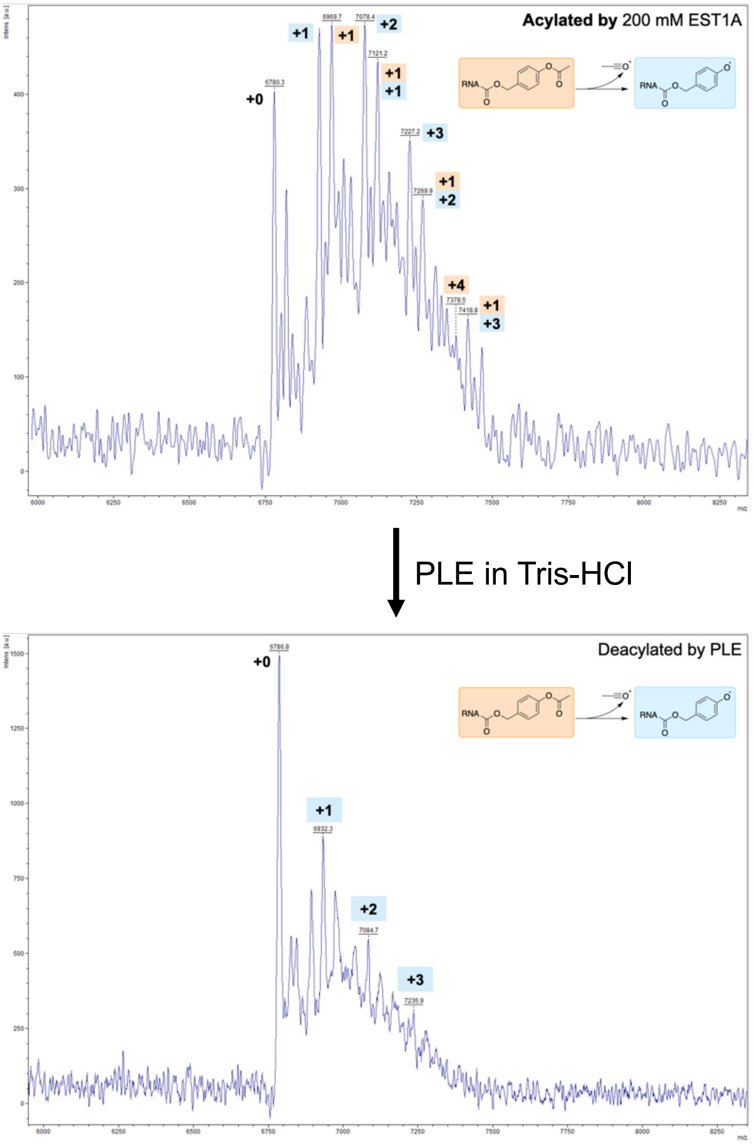


Figure S7. MALDI-TOF spectra of acylated and deacylated siEGFP-AS. Acylation: 5 µM siEGFP-AS incubated with 20 mM **EST1A** in 10% DMSO at 37 for 4 hours. Deacylation: 5 µM acylated siEGFP-AS incubated with 40 U/mL porcine liver esterase (PLE) in 50 mM Tris-HCl (pH 8.0) at 37 °C for 4 hours.

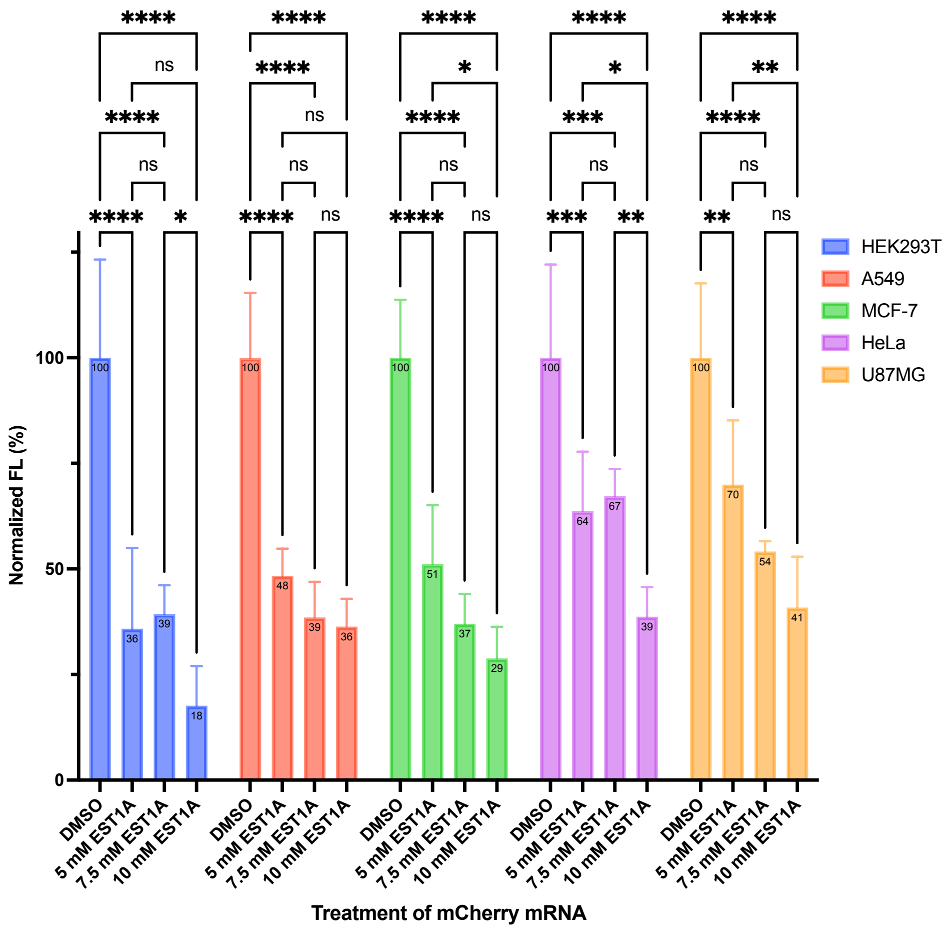


Figure S8. Quantitative analysis of mCheery mRNA functional restoration in different cell lines. Data are shown as mean ± SD (n = 4). t-test: ns, $p > 0.05$; *$p \leq 0.05$; **$p \leq 0.01$; ***$p \leq 0.001$; ****$p \leq 0.0001$.

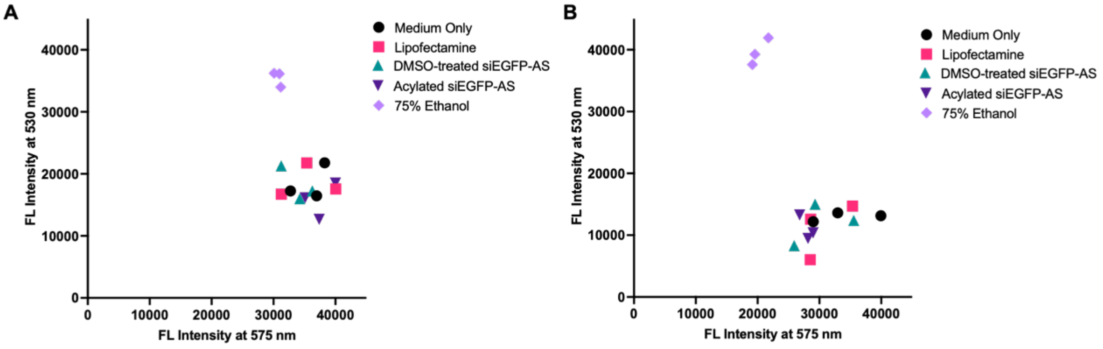


Figure S9. Viability of HepG2 (A) and U87MG (B) cells assessed using the LIVE/DEAD Cell Vitality Kit. For dead-cell controls, cells were treated with 100 µL/well 75% EtOH at 37 °C for 20 min. Graphs show individual data points. Experiments were repeated three times.

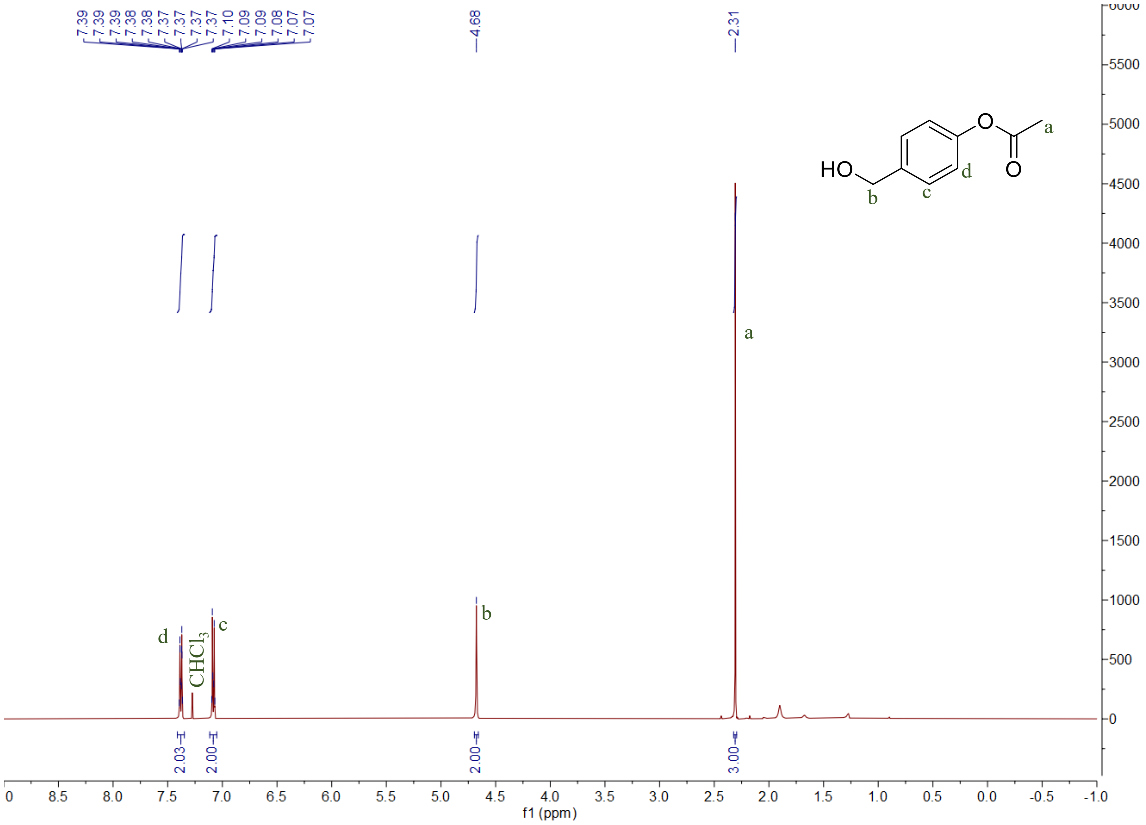
7.39
7.39
7.39
7.38
7.38
7.37
7.37
7.37
7.37
7.10
7.09
7.09
7.08
7.07
7.07
4.68
2.31
HO
O
O
a
b
c
d
CHCl3
2.03
2.00
2.00
3.00
f1 (ppm)

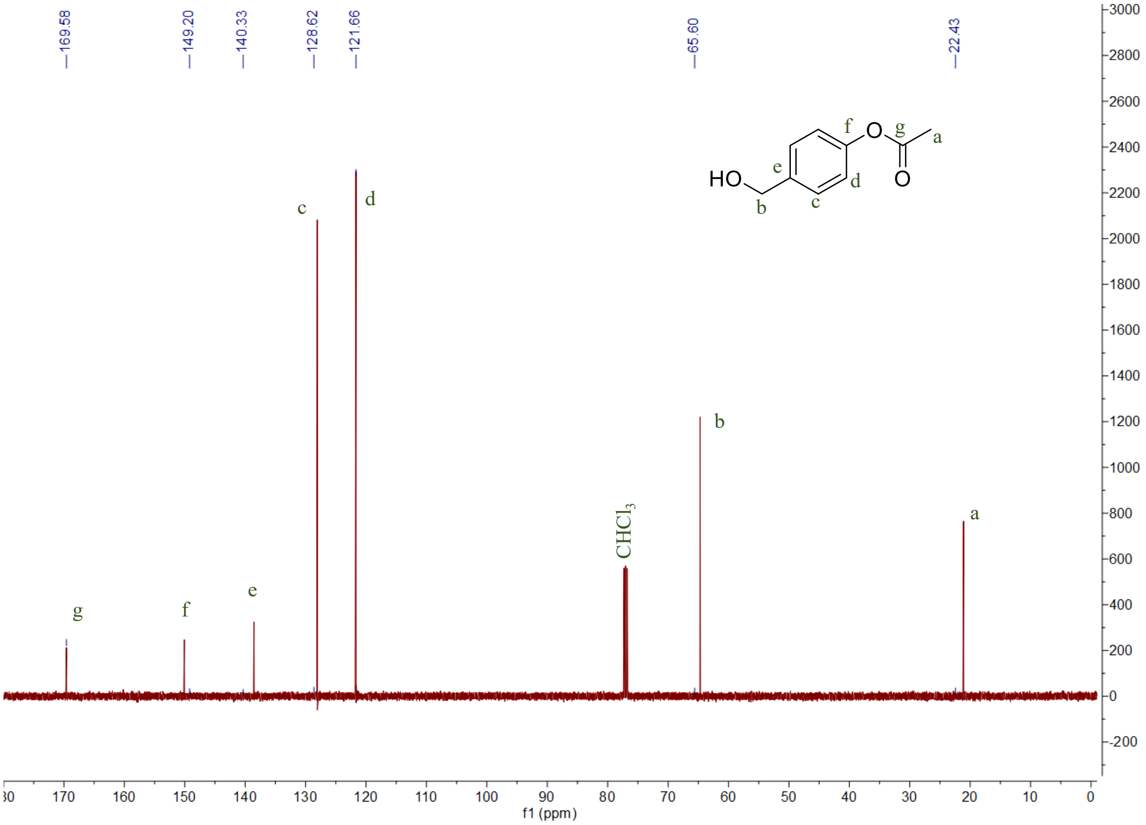
169.58
149.20
140.33
128.62
121.66
65.60
22.43
HO
O
O
a
b
c
d
e
f
g
CHCl3
f1 (ppm)

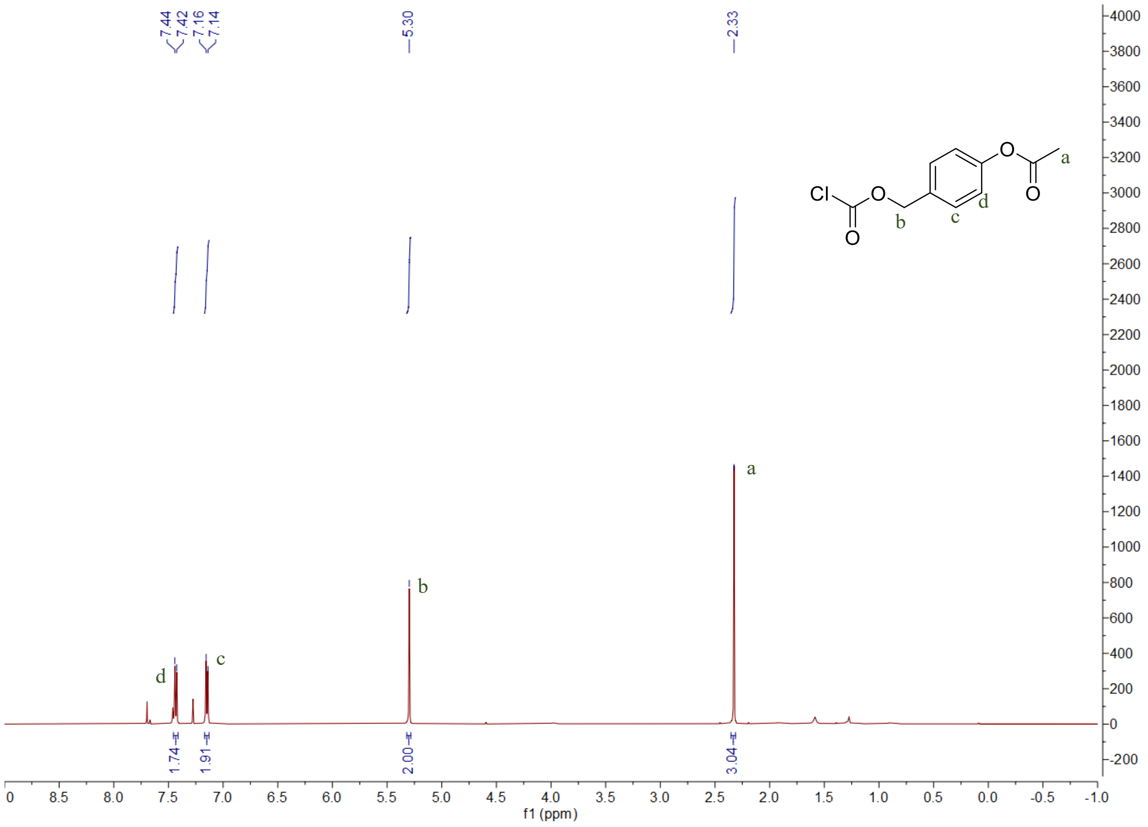
7.44
7.42
7.16
7.14
5.30
2.33
Cl
O
O
O
O
a
b
c
d
1.74
1.91
2.00
3.04
f1 (ppm)

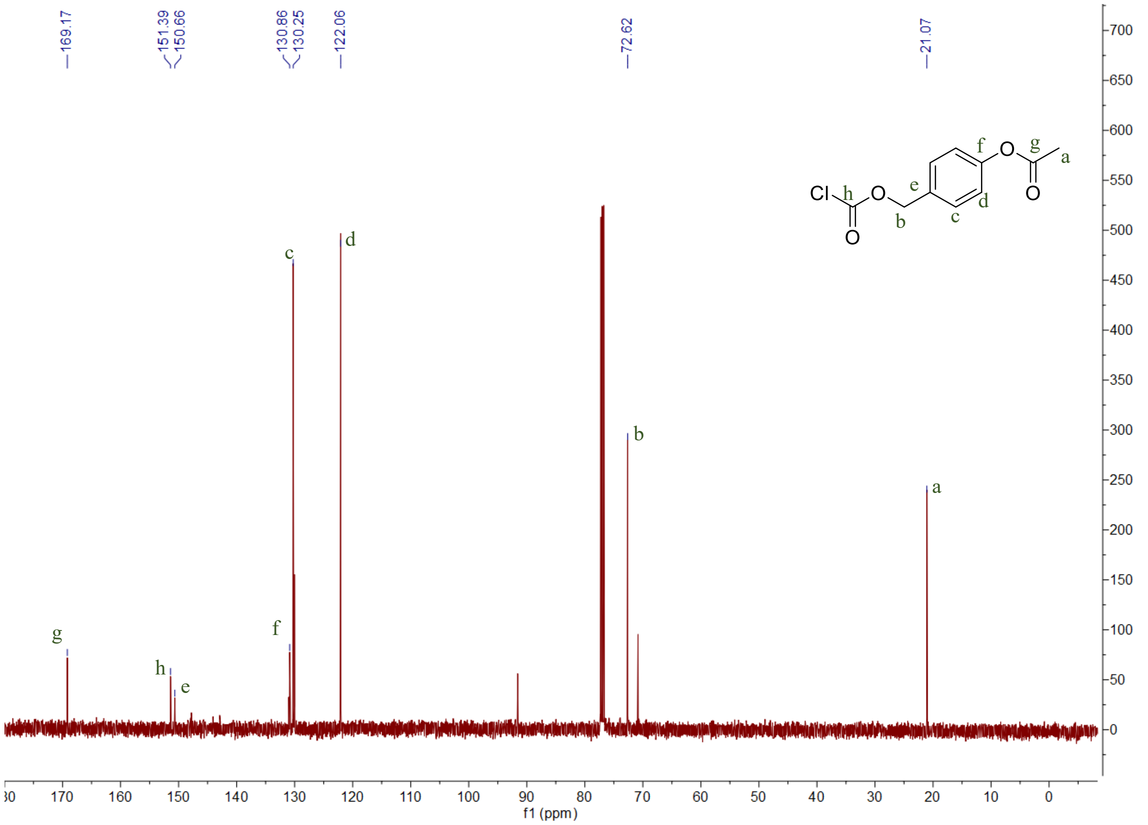
169.17
151.39
150.66
130.86
130.25
122.06
72.62
21.07
Cl
O
O
O
O
a
b
c
d
e
f
g
h
f1 (ppm)

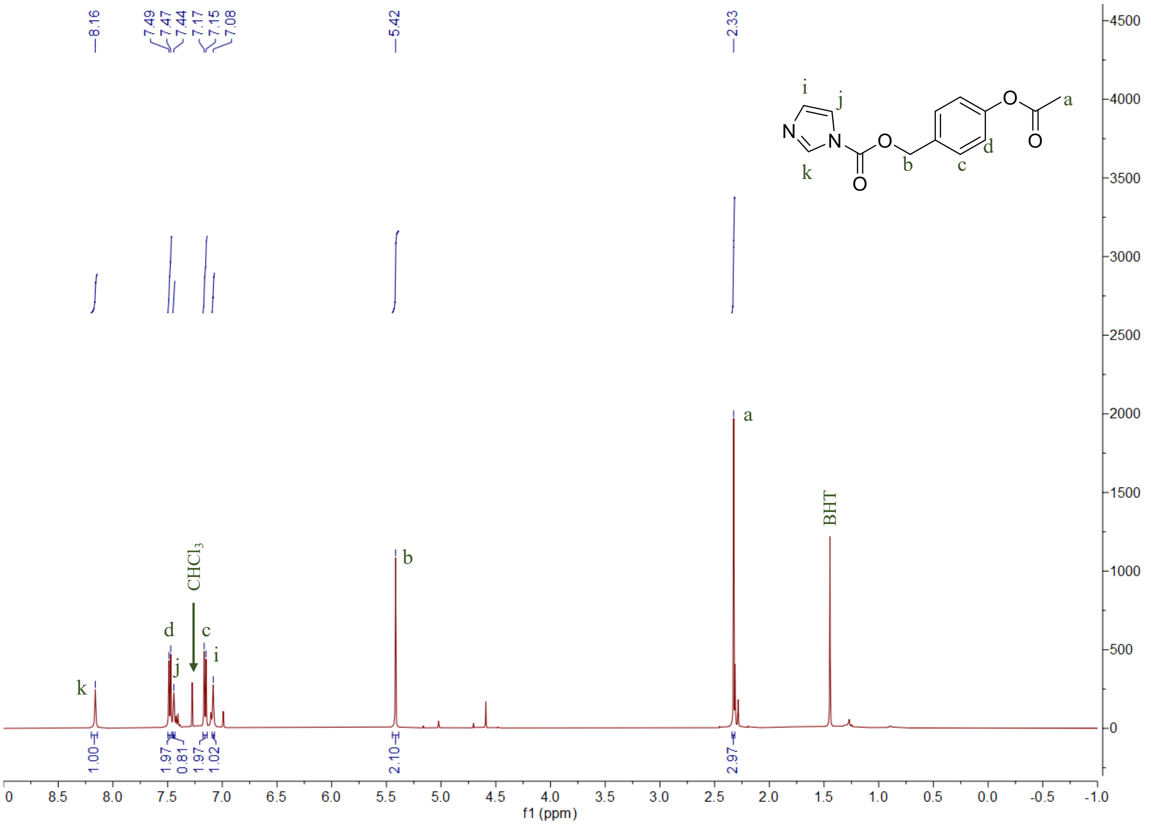

8.16
7.49
7.47
7.44
7.17
7.15
7.08
5.42
2.33
a
b
c
d
i
j
k
N
O
CHCl3
BHT
1.00
1.97
0.81
1.97
1.02
2.10
2.97
f1 (ppm)


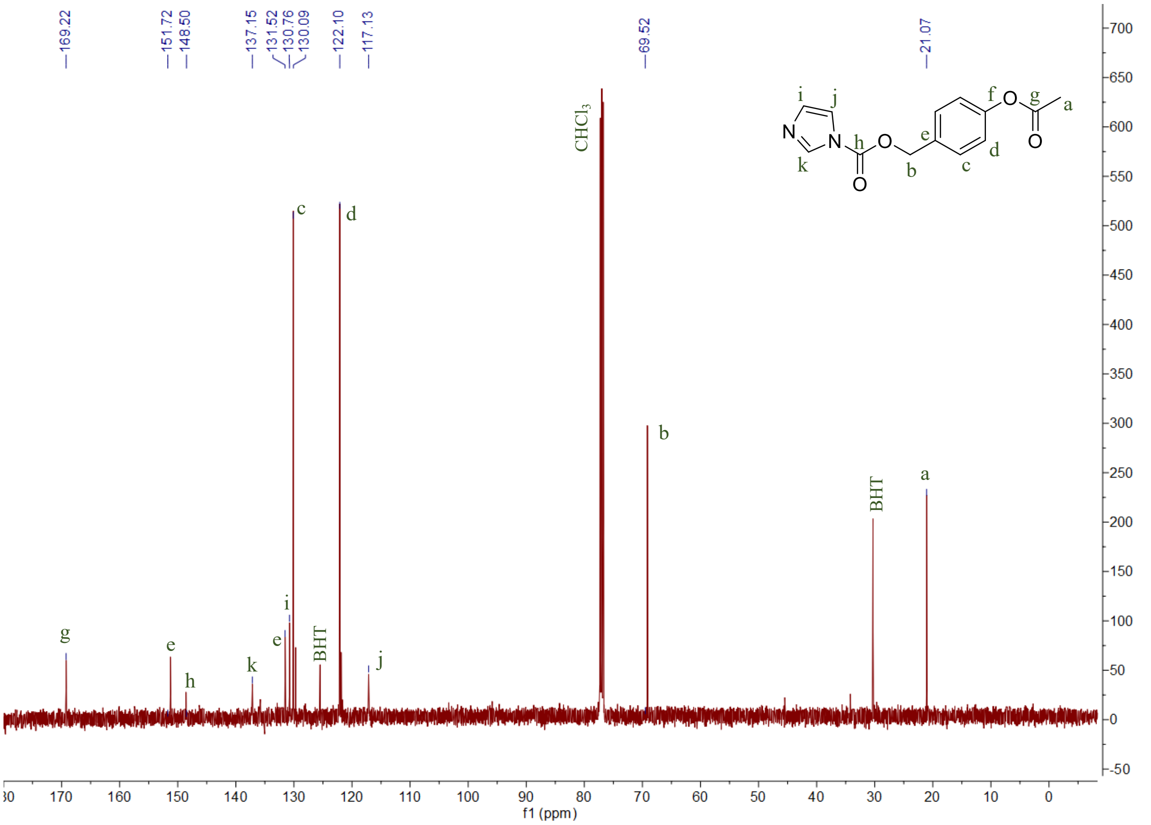

169.22
151.72
148.50
137.15
131.52
130.76
130.09
122.10
117.13
69.52
21.07
a
b
c
d
e
f
g
h
i
j
k
N
O
CHCl3
BHT
f1 (ppm)

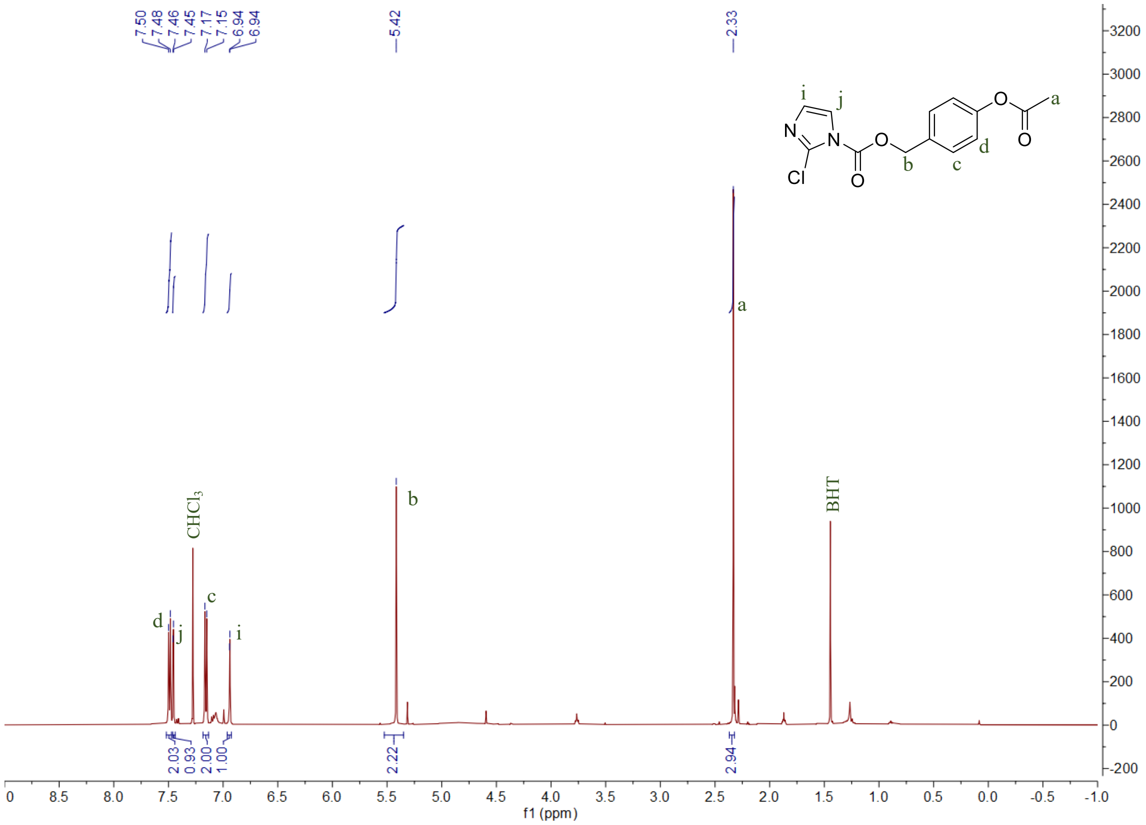
7.50
7.48
7.46
7.45
7.17
7.15
6.94
6.94
5.42
2.33
CHCl3
BHT
f1 (ppm)

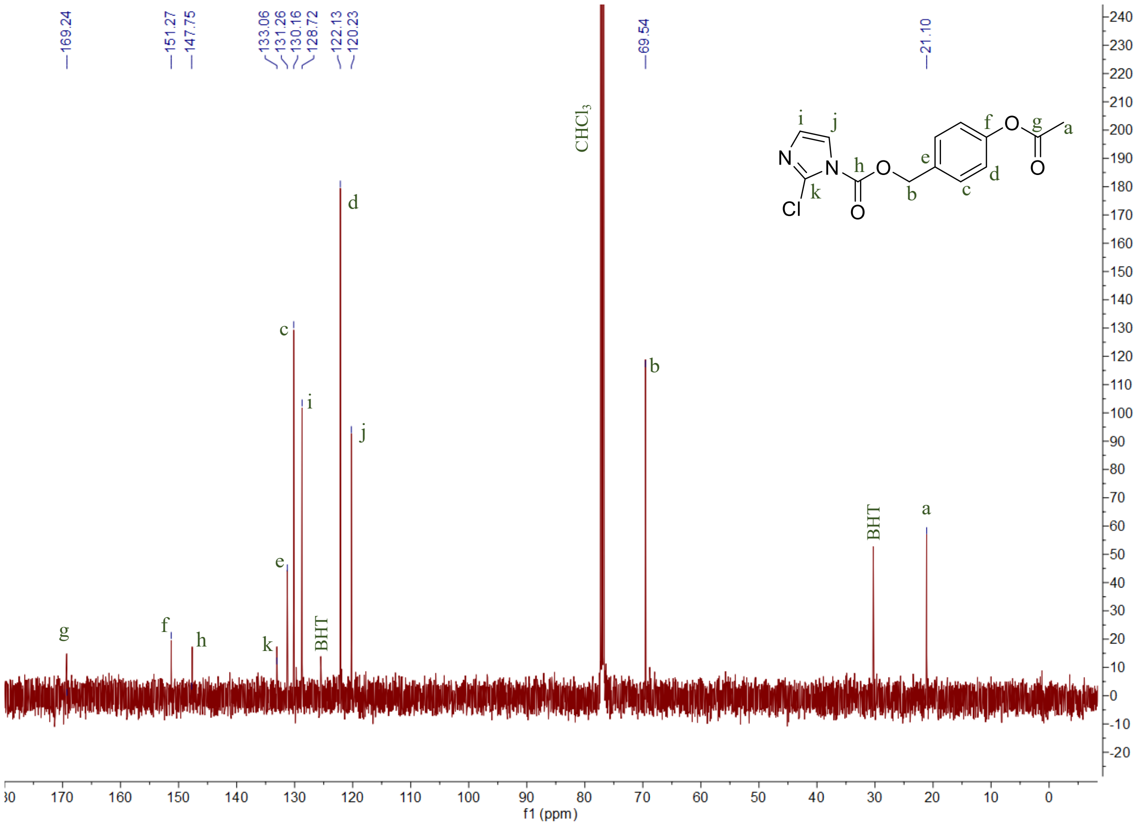
169.24
151.27
147.75
133.06
131.26
130.16
128.72
122.13
120.23
69.54
21.10
CHCl3
BHT
f1 (ppm)

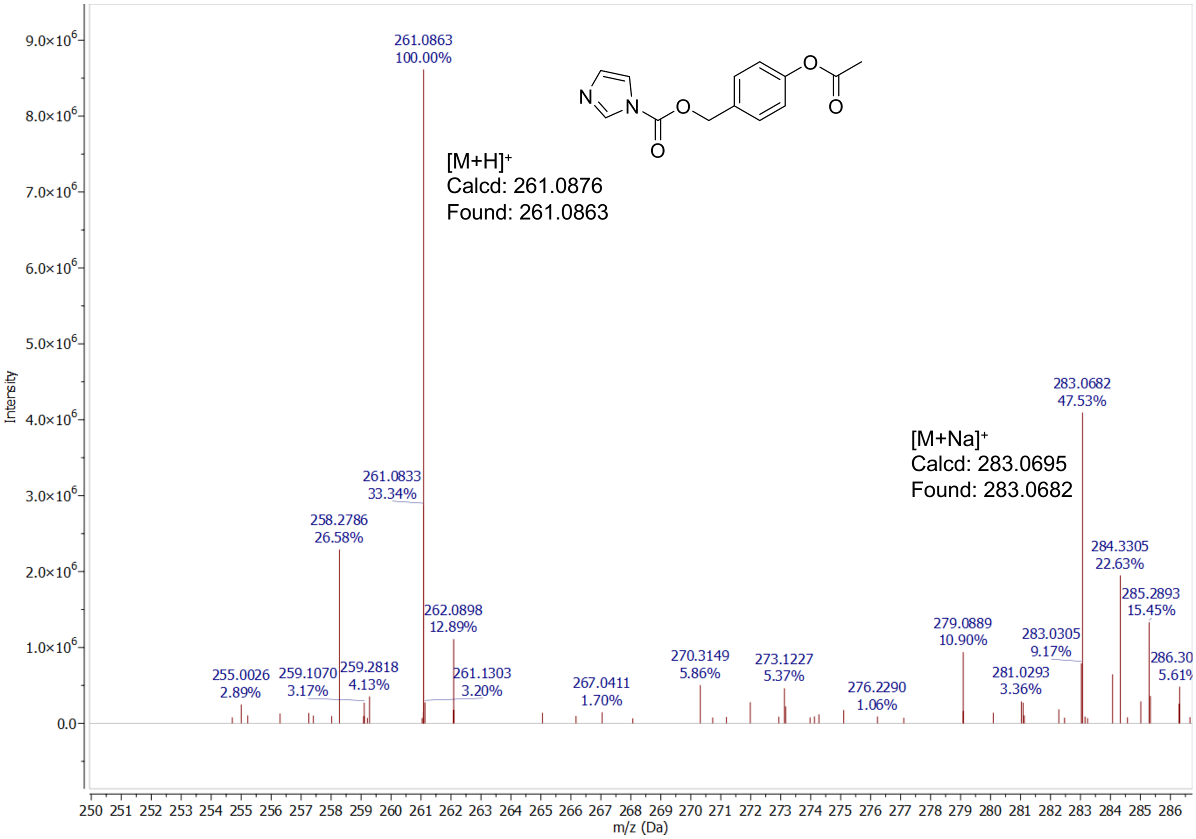
[M+H]+
Calcd: 261.0876
Found: 261.0863
[M+Na]+
Calcd: 283.0695
Found: 283.0682
261.0863
100.00%
283.0682
47.53%
Intensity
m/z (Da)

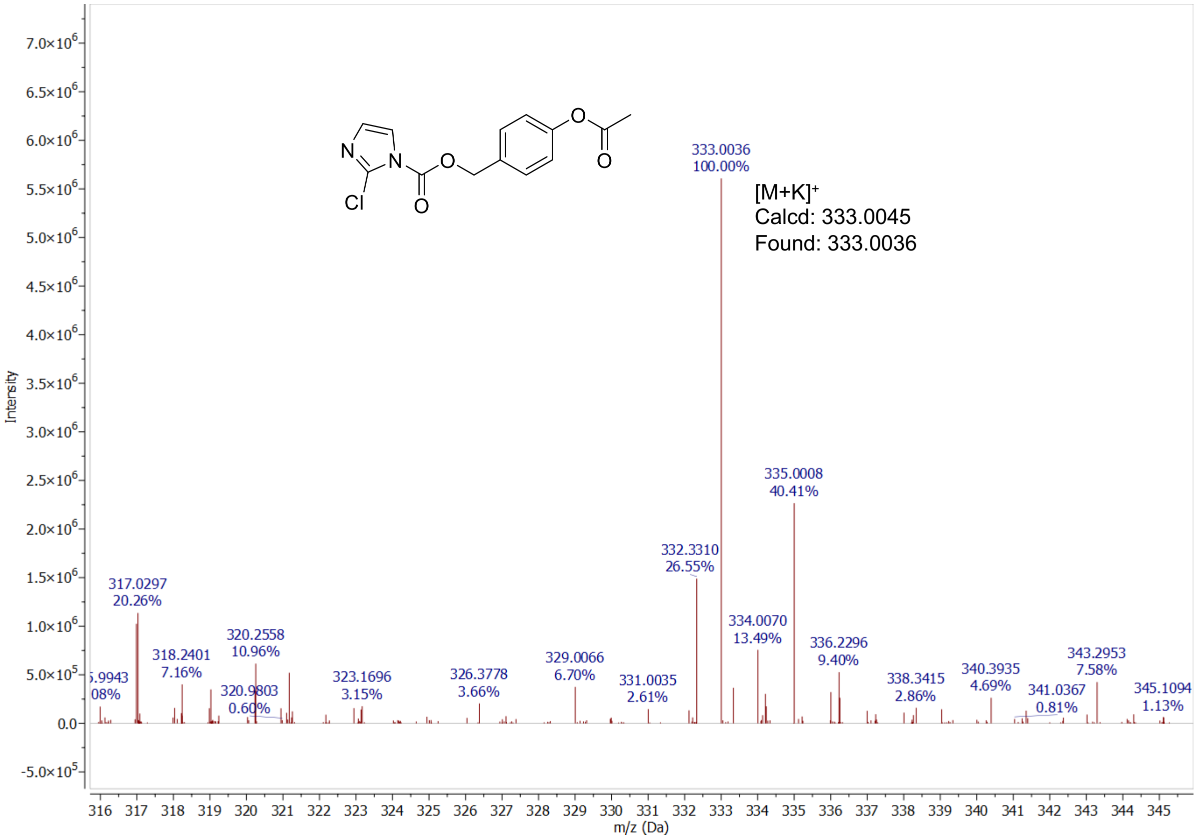
[M+K]+
Calcd: 333.0045
Found: 333.0036
333.0036
100.00%
335.0008
40.41%
Intensity
m/z (Da)

[1] aW. A. Velema, H. S. Park, A. Kadina, L. Orbai, E. T. Kool, *Angew Chem Int Ed Engl* **2020**, *59*, 22017-22022; bK. Liu, R. K. Sriram, M. Rachwalak, W. Yang, A. M. Kietrys, *Cell Reports Physical Science* **2025**, 102928.
